# Plastic germination, temporal niche partitioning, and emergent assortative mating in annual plants

**DOI:** 10.1101/2025.08.15.670279

**Authors:** Max Schmid, Katja Tielbörger, Amaël Daval, Charles Mullon

**Affiliations:** Institute of Evolution and Ecology, University of Tübingen, 72076 Tübingen, Germany; Department of Ecology and Evolution, University of Lausanne, 1015 Lausanne, Switzerland

**Keywords:** predictive germination, phenotypic plasticity, ecological speciation, storage effect, annual plants, genetic linkage

## Abstract

Temporal fluctuations in environmental conditions can promote coexistence via the storage effect, which requires a mechanism buffering different variants during adverse periods. This mechanism readily occurs in annual plants whose seeds do not necessarily germinate each year but instead remain dormant in a seed bank. Yet, how plasticity in germination timing affects genetic diversification and ecological speciation under temporally varying conditions remains poorly understood. Here, we use mathematical modeling and individual-based simulations to investigate the joint evolution of a combination of traits controlling plastic seed germination and plant fecundity under interannual environmental fluctuations. We show that adaptive plasticity in germination readily evolves via a genetic association between seed and plant traits, allowing seeds to germinate preferentially in years favorable for their subsequent growth and reproduction. Such adaptive plasticity enhances the storage effect and promotes genetic divergence among different morphs specialized to distinct yearly conditions. Because these morphs germinate preferentially in different years, plastic germination indirectly generates temporal assortative mating, thereby maintaining genetic associations among seed and plant traits despite the absence of physical linkage between loci controlling these traits. This temporal assortative mating ultimately leads to reproductive isolation between morphs, thus laying the foundation for ecological speciation. Our findings show that adaptive plasticity and genetic diversification are not mutually exclusive but can interact synergistically to enhance biodiversity.

## 1 Introduction

Most communities experience fluctuations in environmental conditions, with year-to-year variation in temperature, rainfall, or disturbances such as fires or floods (Pausas and Lamont, 2022; He et al., 2019). Phenotypic adaptation to these interannual fluctuations can occur in two main ways. First, individuals can respond plastically to prevailing conditions, resulting in environmentally induced phenotypes (polyphenism; Woltereck 1909; Bradshaw 1965; Scheiner 1993; Schlichting 1986; Via et al. 1995). The evolution of adaptive plasticity requires reliable cues that predict the selective environment (Moran, 1992; Tufto, 2000; Bonamour et al., 2019), as well as sufficiently flexible developmental programs (DeWitt et al., 1998; Lande, 2014; Dupont et al., 2024). Second, populations can evolve genetic polymorphisms, or species within communities can diverge via character displacement, leading to temporal niche partitioning. In both cases, different variants (alleles or species) specialize on contrasting yearly conditions (Chesson and Warner, 1981; Ellner and Hairston, 1994; Yamamichi et al., 2023). The evolution of temporal niche partitioning requires a mechanism buffering variants against selection in unfavorable years, known as the temporal storage effect (Chesson, 1983; Svardal et al., 2015; Yamamichi et al., 2023). Determining how these processes shape genetic and phenotypic variation is essential for understanding biodiversity and its maintenance in temporally variable environments.

Adaptive plasticity and genetic polymorphism are typically viewed as alternatives but these processes may also interact (West-Eberhard, 1989; Crispo, 2008; Conover et al., 2009; Ghalambor et al., 2015; Schmid and Guillaume, 2017; Sommer, 2020). One context where this interaction might be particularly relevant is temporal niche partitioning in annual plants. Under interannual fluctuations, niche partitioning readily evolves when seeds exhibit between-year dormancy, whereby a fraction remains dormant for multiple years before germinating (Chesson, 1994; Ellner and Hairston, 1994; Chesson, 2000; Barabás et al., 2018; Johnson and Hastings, 2022), generating the necessary storage effect. Seed germination itself can show adaptive plasticity, with seeds germinating preferentially under conditions predicting favorable subsequent growth and reproduction (e.g., Angert et al., 2009; Donohue et al., 2010; Cohen, 1967). Such predictive germination (Venable and Lawlor, 1980; Pake and Venable, 1996) can reinforce character displacement, causing different species to germinate in different years, thereby facilitating coexistence via the storage effect (Kortessis and Chesson, 2021). However, previous studies assumed divergence among already differentiated species (Venable et al., 1993; Snyder and Adler, 2011; Kortessis and Chesson, 2021), leaving unresolved whether predictive germination can initiate genetic diversification and ecological speciation within initially homogeneous populations.

Here, we investigate whether the evolution of predictive germination facilitates the emergence and divergence of genetic morphs within annual plant species. Using mathematical modeling and individual-based simulations, we show that predictive germination enhances the conditions under which temporal niche partitioning evolves. We further explore evolutionary constraints imposed by sexual reproduction and recombination, revealing that adaptive divergence in germination timing generates temporal assortative mating, thus setting the stage for ecological speciation.

## 2 Model

### 2.1 Life cycle and traits

We model a large population of annual plants that shows between-year seed dormancy (though this model can equally apply to other taxa, such as *Daphnia* with resting eggs or fungi with dormant spores). Seeds may remain dormant in the seed bank for multiple years, while plants that germinate are short-lived and die after reproduction.

The population may experience environmental conditions during germination and reproduction that vary between years (e.g., fluctuations in humidity, temperature, or herbivore density). We represent these fluctuations with two random variables per year *t* : *θ*_g,*t*_, describing the environment during the germination period, and *θ*_f,*t*_, describing the environment later in the year during reproduction. These conditions influence germination and reproduction through interactions with specific plant traits, as detailed below.

#### 2.1.1 Seed germination

At the beginning of each growing season, seeds in the seed bank either remain dormant or germinate. The probability that a seed indexed *i*, characterized by its germination trait *z*_g,*i*_, germinates in year *t* depends on the early-year environment (*θ*_g,*t*_) as follows:

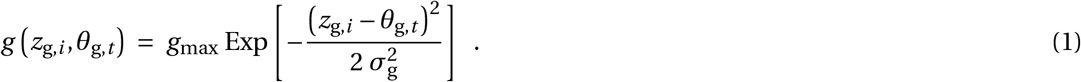

Equation (1) indicates that a seed germinates with a maximum probability, 0 *< g*_max_ *≤* 1, when its trait matches environmental conditions (i.e., when *θ*_g,*t*_ *= z*_g,*i*_ ; Fig. 1C). In this view, trait *z*_g,*i*_ could be seen as a physiological or morphological seed character that regulates the preferred environment for germination. Germination probability declines with increasing mismatch between a seed’s trait and the environment, at a rate determined by 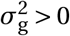.

**Figure 1.**
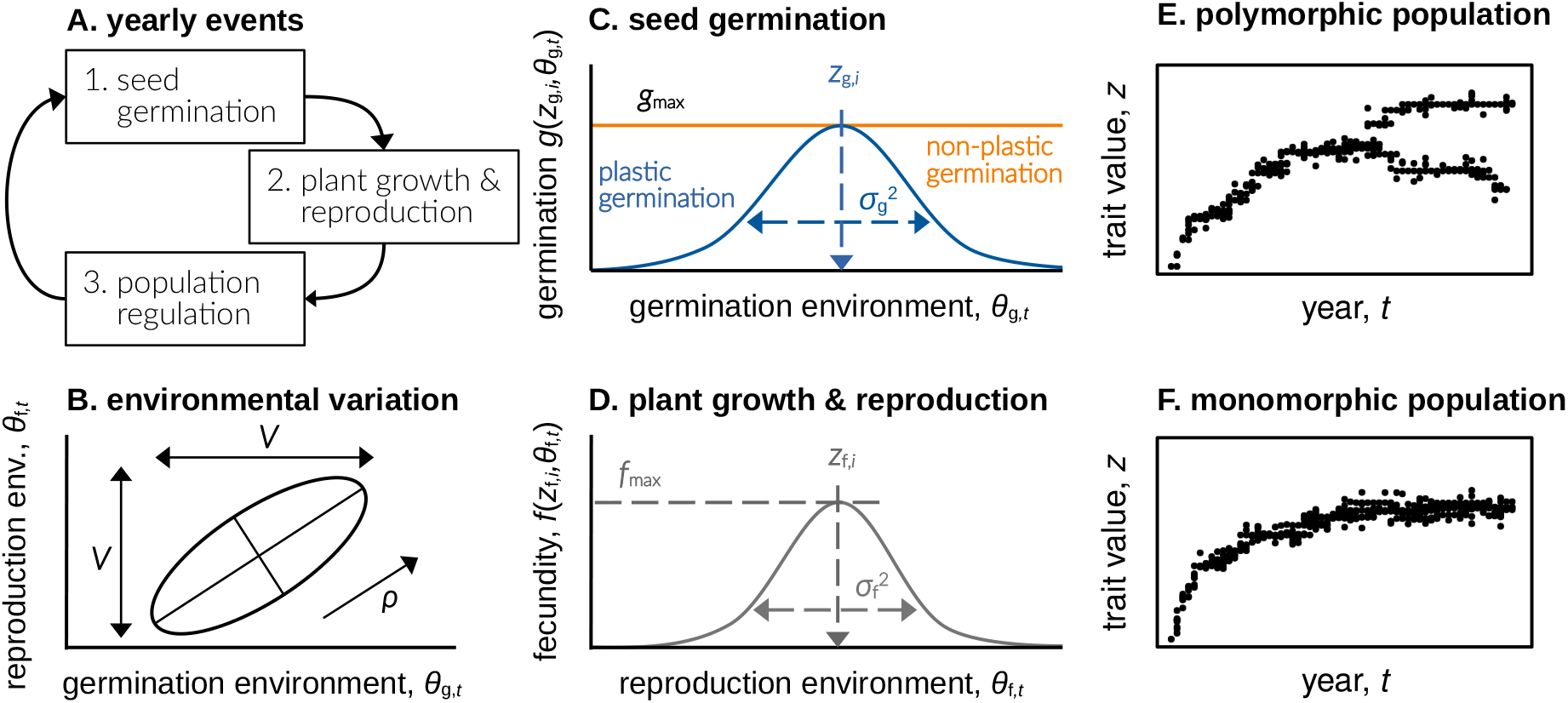
Graph **A** shows the yearly life-cycle events, where a fraction of seeds germinates, reproduces, and the plant population then experiences density regulation. The ellipsoid in graph **B** illustrates the environmental distribution where environmental conditions during germination *θ*_g,*t*_ (x-axis) and reproduction *θ*_f,*t*_ (y-axis) are drawn from a bivariate normal distribution, each year anew. Both environmental conditions show between-year variance *V* and within-year environmental correlation *ρ*. Graph **C** contrasts plastic seed germination (blue line) with non-plastic seed germination (orange line, where 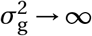). Graph **D** shows how the individual fecundity changes with the match between *θ*_f,*t*_ and the individual fecundity trait *z*_f,*i*_. The evolutionary dynamics in *z*_g_ and *z*_f_ then could either lead to a polymorphic plant population under disruptive selection (**E**), or to a monomorphic population with stabilizing selection (**F**).

Equation (1) sets a reaction norm describing how seed germination responds to the environment with the parameter 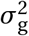 tuning the degree of germination plasticity. A small 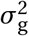 corresponds to high plasticity, meaning germination probabilities change strongly in response to environmental variation (blue line in Fig. 1C; e.g., see Simons, 2014, for empirical data supporting bell-shaped germination functions). In contrast, large 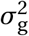 corresponds to low plasticity, meaning germination probabilities remain relatively constant across a broad range of environmental conditions. In the limiting case 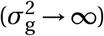, germination becomes completely independent of the environment (non-plastic), reducing the reaction norm to a horizontal line with intercept *g*_max_ (orange line in Fig. 1C; e.g., see Ellner and Hairston, 1994).

#### 2.1.2 Plant growth and reproduction

After germination, plants grow and reproduce. We assume that the fecundity of plant *i*, characterized by its fecundity trait *z*_f,*i*_, in response to the reproduction environment *θ*_f,*t*_, is given by

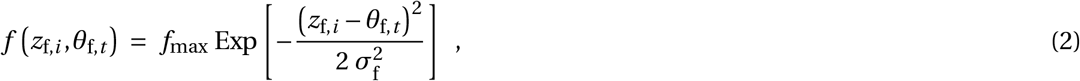

where maximum fecundity *f*_max_ is reached when the fecundity trait exactly matches environmental conditions during reproduction (i.e., when *z*_f,*i*_ *= θ*_f,*t*_ ; Fig. 1D). Trait *z*_f,*i*_ could be seen as a quantitative trait in germinated plants that governs the preferred environment for reproduction (like root morphology or stomata density). Fecundity declines with increasing mismatch between trait and environment at a rate determined by 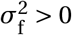. This parameter 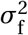 measures the breadth of the fecundity niche, with large 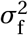 indicating a broad niche (plants achieve high fecundity across a wide range of conditions) and small 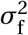 indicating a narrow niche (high fecundity is limited to specific environmental conditions). Eq. (2) represents a classical assumption in many theoretical models of adaptation to environmental variables (e.g., Via and Lande, 1985; Day, 2000; Kisdi, 2002; Chevin and Lande, 2011; Svardal et al., 2015; Miller and Klausmeier, 2017; Yamamichi et al., 2019; Ohtsuki et al., 2020; Orive et al., 2023; Saltini et al., 2023).

#### 2.1.3 Regulation

After reproduction, all germinated plants die. Seeds that did not germinate remain dormant, surviving to the next year with probability *s* and dying with probability 1*−s*. Newly produced seeds compete for the available spots in the seed bank, which become available when seeds germinate or die. We assume the total number of seeds in the seed bank remains constant over time such that *N* seeds are present at the start of each year (with *N* large). Competition for these available seed-bank spots is random with respect to trait values as per the lottery model (Chesson and Warner, 1981).

### 2.2 Environmental heterogeneity and genetic constraints

We assume environmental conditions during germination (*θ*_g,*t*_) and reproduction (*θ*_f,*t*_) fluctuate randomly among years around the same long-term mean and with equal variance 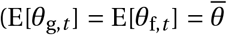 and V[*θ*_g,*t*_] *=* V[*θ*_f,*t*_] = *V* for all *t* ; Supplementary Appendix A). Conditions during germination and reproduction within the same year may be correlated (Corr[*θ*_g,*t*_, *θ*_f,*t*_] = *ρ* for all *t* ; Fig. 1B; e.g., see Kortessis and Chesson 2021). A positive within-year correlation (*ρ >* 0) implies predictive germination cues (e.g., dry germination periods predict dry reproductive periods), whereas a negative correlation (*ρ <* 0) indicates an inverse relationship (e.g., dry germination periods predict wet reproductive periods). Weak correlations (*ρ ≈* 0) imply limited predictability between conditions at germination and reproduction, likely impeding the evolution of predictive germination.

We explore how genetic constraints interact with environmental correlations to influence adaptation and diversification. We consider the joint evolution of the germination (*z*_g_) and fecundity (*z*_f_) trait, which together determine environmental conditions at germination and subsequent reproduction. We compare two scenarios: (i) the *single-trait case*, where germination and fecundity traits are genetically constrained to be identical (*z*_g,*i*_ *= z*_f,*i*_ ; such that each individual maximizes germination and subsequent fecundity at the same condition), and (ii) the *two-trait case*, where traits evolve independently (such that the each individual may prefer different conditions for germination and for reproduction). Comparing these scenarios allows examination of how genetic constraints influence the evolution of temporal niche partitioning under different degrees of environmental correlation.

### 2.3 Analyses

We aim to identify the ecological and genetic conditions under which germination and fecundity traits become polymorphic, thus leading to temporal niche partitioning. We use two complementary approaches.

First, we conduct evolutionary invasion analyses (Supplementary Appendices B-C), assuming haploid, clonally reproducing plants. Each trait is encoded by a single locus evolving under the continuum-of-alleles model. Mutations are rare and have small phenotypic effects such that evolution occurs in two phases (Geritz et al., 1998). Initially, directional selection drives the population toward convergence-stable trait values with the population remaining largely monomorphic. Subsequently, the population either remains monomorphic under stabilizing selection (Fig. 1E) or undergoes evolutionary branching, becoming genetically and phenotypically polymorphic under disruptive selection (Fig. 1F). Our goal is to identify conditions favoring evolutionary branching.

Second, we use individual-based simulations, relaxing key analytical assumptions by explicitly incorporating diploidy, sexual reproduction, and varying genetic linkage between germination and fecundity loci. These simulations test the robustness of our analytical predictions and allow us to investigate how sexual reproduction and recombination influence the evolution of trait polymorphism and temporal niche partitioning (Supplementary Appendix D).

## 3 Results

### 3.1 Germination plasticity facilitates polymorphism under positive environmental correlations and genetic constraints

We first investigate the *single-trait case* where individuals achieve maximum germination and maximum reproduction at the same environmental condition (i.e. *z*_g,*i*_ *= z*_f,*i*_ due to genetic constraints). As a baseline, we compare our results to the classical scenario without plasticity (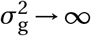; Fig. 1C), for which polymorphism evolves under the condition 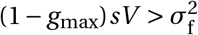 (orange areas in Fig. 2; Chesson, 1983; Ellner and Hairston, 1994; Svardal et al., 2015). Polymorphism in this baseline scenario is thus favored by high between-year environmental variability (*V*), narrow fecundity niches (small 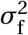), and extensive seed dormancy (small *g*_max_, large *s*).

**Figure 2.**
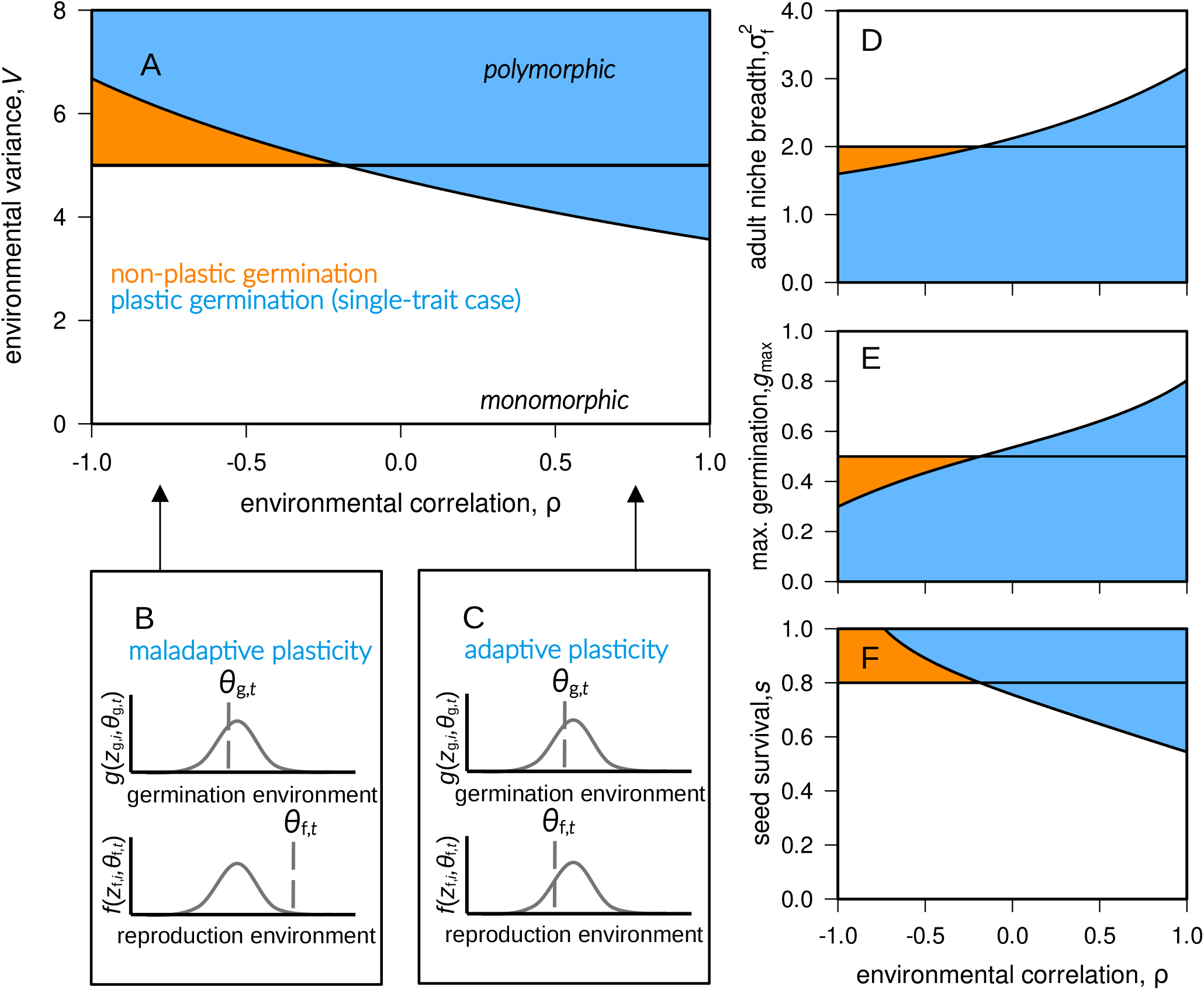
Graphs **A, D**-**F** present analytical results for the *single-trait case* (*z*_g,*i*_ =*z*_f,*i*_) where plants evolve genetic/phenotypic polymorphism in the colored areas, but stay monomorphic in the white areas. The case of plastic seed germination is presented in blue (when 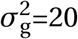; according to eq. B13 with components B16), the case of non-plastic germination is shown in orange (when 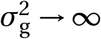; following 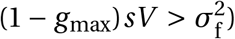. If not specified in graph B and D-F, the parameter values are: 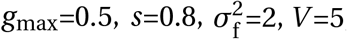, and *ρ*=1. Graphs **B** and **C** conceptualize how environmental correlation leads to adaptive and maladaptive germination plasticity. With *ρ <* 0 and large differences between *θ*_g,*t*_ and *θ*_f,*t*_ within years, a specialized plant morph 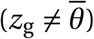 might experience suitable conditions for germination, but detrimental conditions during reproduction (graph B). With *ρ >* 0, environmental conditions do not change much within years (*θ*_g,*t*_ *≈ θ*_f,*t*_) such that plants with *z*_g,*i*_ *= z*_f,*i*_ could combine high germination probabilities with high fecundities.

When allowing plastic germination 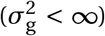, conditions for evolutionary branching are more involved mathematically (see eq. B16 in Appendix B) but can be easily interpreted graphically (Fig. 2). Plastic germination broadens conditions leading to polymorphism under positive within-year environmental correlations (blue areas in Fig. 2, *ρ >* 0). Here, plasticity allows plants to germinate preferentially in years reliably favorable for subsequent reproduction; that is, plasticity in this case is adaptive (following Gotthard and Nylin, 1995‘s conceptual distinction). For instance, plasticity evolves such that plants adapted to warm conditions germinate and reproduce predominantly in warm years, while those adapted to cooler conditions germinate predominantly in cooler years. By allowing plants specialized to distinct environmental conditions to reproduce mostly in separate years, plastic germination here reduces between-morph competition and enhances temporal niche partitioning.

Conversely, plastic germination restricts polymorphism relative to the non-plastic scenario under negative environmental correlations (blue areas in Fig. 2, *ρ <* 0). In this situation, the same environmental preference during germination and fecundity cannot simultaneously match contrasting environmental conditions within a year (i.e., *θ*_g,*t*_*≠ θ*_f,*t*_ when *ρ → −*1; see eq. A10). This mismatch means seeds germinate into conditions detrimental for reproduction, so germination plasticity here is maladaptive (Fig. 2B). For example, warm-adapted plants germinate preferentially under warm conditions but then experience cold conditions during reproduction. Such mismatch in turn hinders temporal niche partitioning.

Overall, we see that adaptive germination plasticity promotes polymorphism while maladaptive plas-ticity opposes it. This parallels findings by Kortessis and Chesson (2021), who showed that predictive germination promotes divergence between differentiated species when reliable cues are present and when the germination schedule can be aligned to beneficial conditions for plant growth.

In the absence of environmental correlation within years (*ρ =* 0), germination plasticity can either facilitate or oppose polymorphism relative to the non-plastic scenario, depending on parameters (Fig. 2, horizontal line at *ρ =* 0; eq. B18). Two opposing effects explain this result. On the one hand, increased plasticity (smaller 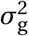) reduces the long-term average germination probability, increasing seed dormancy and thus strengthening the storage effect, thereby favoring polymorphism. On the other hand, increased plasticity induces stabilizing selection on the trait during germination when environmental variability (*V*) or seed survival (*s*) is limited (in fact, selection is stabilizing if 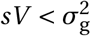 when fecundity selection is weak, 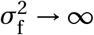). The interplay of these two effects determines whether germination plasticity promotes or inhibits polymorphism when *ρ =* 0.

### 3.2 Temporal niche partitioning under positive and negative environmental correlation when germination and fecundity traits evolve independently

We next allow germination and fecundity traits to evolve independently (*two-trait case*). Allowing independent trait evolution broadens conditions leading to polymorphism, now occurring under both strongly positive and strongly negative environmental correlations within years (high absolute values of *ρ*, dark blue areas in Fig. 3B). Under negative correlation (*ρ <* 0), polymorphism emerges because germination and fecundity traits can evolve in opposite directions, each trait matching distinct environmental conditions within the same year. For example, some plants evolve to germinate preferentially under cool conditions but reproduce best under warm conditions, while others evolve the opposite pattern. This generates negative covariance between germination and fecundity traits (*z*_g,*i*_ and *z*_f,*i*_ ; Fig. 3D, see also eq. C19 in Appendix C). Conversely, under positive environmental correlation (*ρ >* 0), covariance between germination and fecundity traits becomes positive, aligning germination timing with conditions beneficial for reproduction (Fig. 3E). Germination plasticity is thus adaptive at both positive and negative correlation, enhancing temporal niche partitioning whenever environmental correlations within years are sufficiently strong.

**Figure 3.**
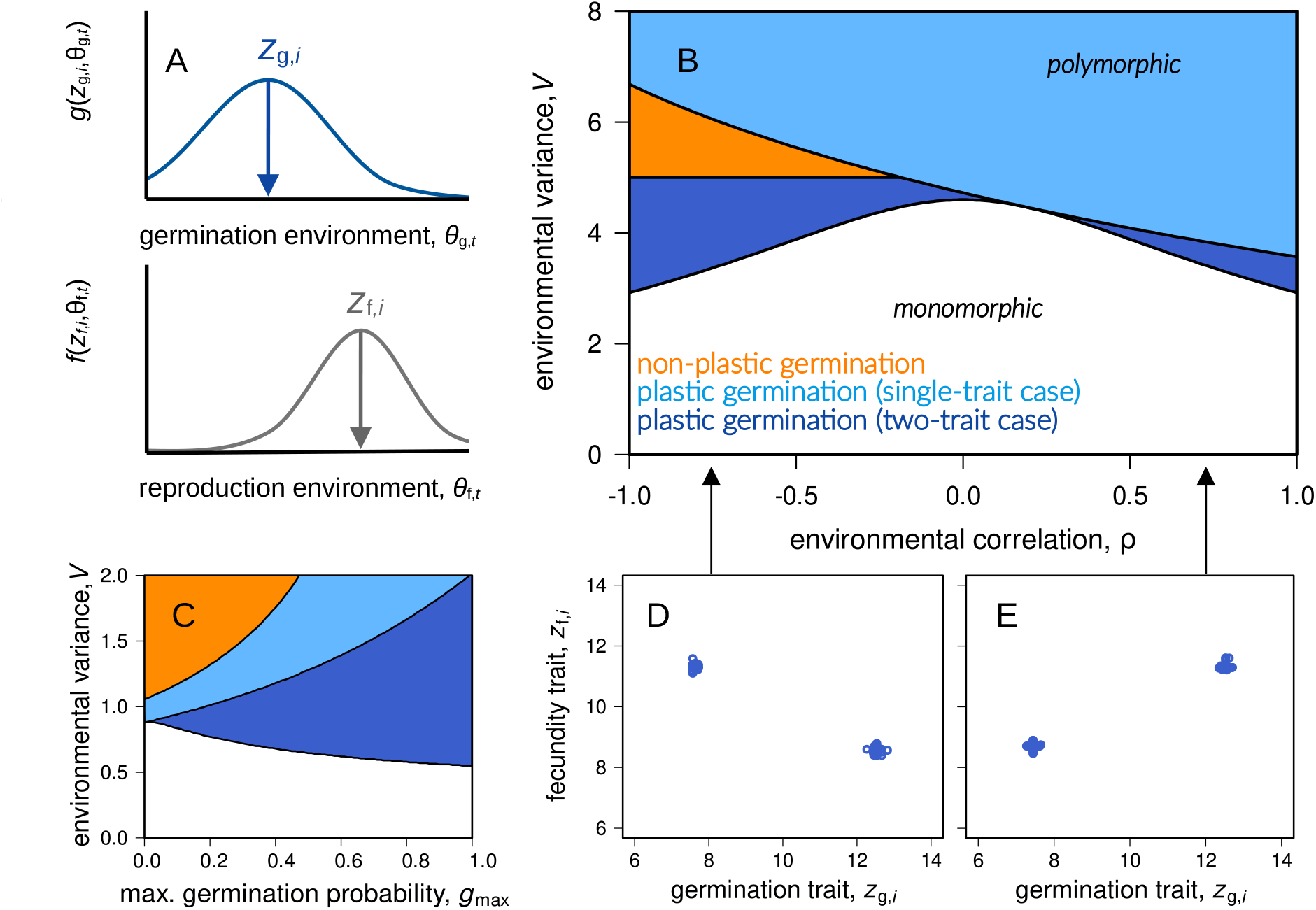
Graph **A** illustrates the *two-trait case* where the germination trait and the fecundity trait could evolve independently and differ from each other within individuals (such that *z*_g,*i*_ ≠ z_f,*i*_). The analytical results for the two-trait case in graphs **B** and **C** present conditions leading to trait poly-morphism and niche partitioning (dark blue area), and conditions leading to monomorphic plant populations (white area; from the Hessian matrix C6 with components C17-C19). Graphs **D**-**E** illustrate how plants could diversify with negative (positive) environmental correlation by the evolution of negative (positive) covariance between individual *z*_g,*i*_ and *z*_f,*i*_ values. Parameter values for graphs B),D),E) are: 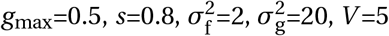, with *ρ*=1 (graph E) and *ρ*=-1 (graph D). The parame-ters for graph C) are: 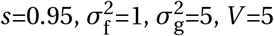, and *ρ*=1.

In the two-trait scenario, high maximum germination probabilities (large *g*_max_) can facilitate polymorphism, particularly under strong environmental correlations (large |*ρ*|) and high seed survival (*s*; dark blue area in Fig. 3C). This contrasts with our single-trait scenario and with earlier models (e.g., Wisnoski and Shoemaker, 2022), where increased dormancy (lower *g*_max_) promotes polymorphism by enhancing the storage effect. This difference arises because independent evolution of the two traits enables the evolution of highly adaptive plasticity when |*ρ*| is large: specialized genotypes evolve to germinate almost exclusively in conditions that reliably predict high fecundity. As a consequence, increasing the maximum germination probability (*g*_max_) allows these specialized genotypes to express their traits more frequently when conditions are favorable. Seed survival during dormancy (*s*) modulates this effect by enabling seeds to persist through unfavorable periods, thus maintaining the specialized genotypes across years and sustaining the storage effect.

### 3.3 Plastic seed germination drives phenotypic variation within and among years

We next examined the phenotypic distributions at mutation-selection-drift equilibrium that arose in the individual-based simulations. We measured phenotypic variance in the germination and fecundity traits (*P*_g,w_ and *P*_f,w_; see eq. D5) in germinated plants. Conditions favoring polymorphism also led to greater phenotypic variance. Specifically, high environmental variability (*V*) and strong environmental correlation (|*ρ*|) resulted in increased phenotypic variance in both germination and fecundity traits (Fig. 4A,F). This occurred because these conditions generated strong selection for coordinated trait combinations that were specialized to distinct yearly environmental conditions. When selection was sufficiently strong, more than two plant morphs could evolve and coexist (Fig. 5D).

**Figure 4.**
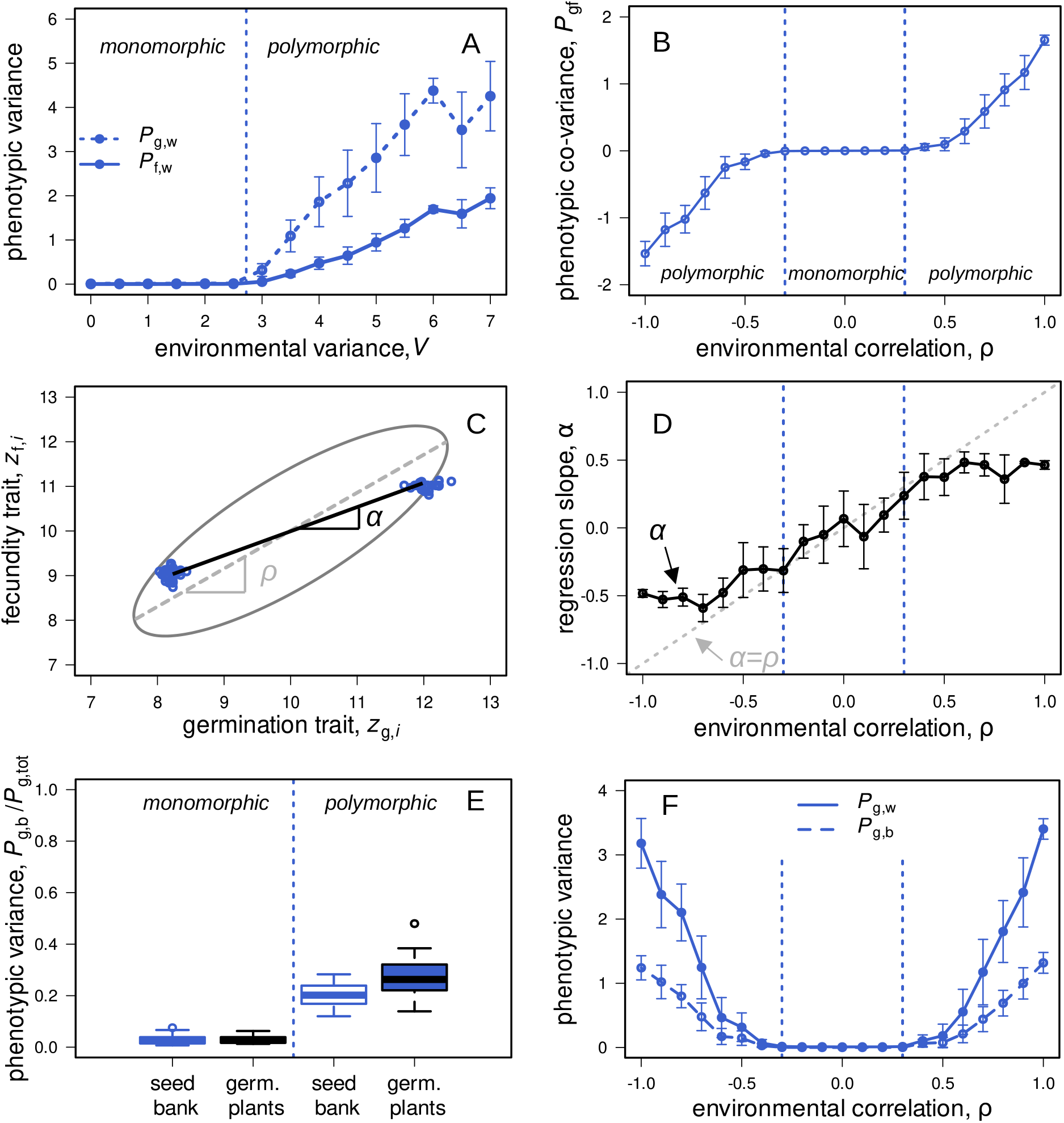
These graphs show simulation results for the two-trait case with clonal reproduction in haploid plants. Graph **A** depicts within-year phenotypic variance in the germination trait (dashed lines) and the fecundity trait (solid lines) as a function of interannual environmental variation *V*. The vertical dotted lines (—) present the threshold between stabilizing and disruptive selection as predicted by the mathematical model (following the Hessian matrix C6 with components C17-C19). Graph **B** illustrates the extent of trait covariance *P*_gf_ (see eq. D7) as a function of environmental correlation, where polymorphism could evolve with positive and negative *ρ*. Graph **C** conceptualizes the linear regression slope for the phenotypic distribution (*α*) and for the environmental distribution (*ρ*), where plants not always evolve *α = ρ* (graph **D**). Graph **E** shows simulation results of the between-year phenotypic variance (*P*_g,b_/*P*_g,tot_) for seeds in the seed bank and for germinated plants only. Graph **F** presents the between-year phenotypic variance as a function of environmental correlation. The parameter values for graphs A-F are: 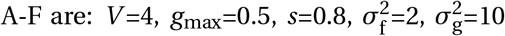 (with *ρ*=0.8 for graph C; and *ρ*=(0,1) for graph E). The whiskers in graphs A,B,D,F illustrate variation among replicates (covering one standard deviation).

**Figure 5.**
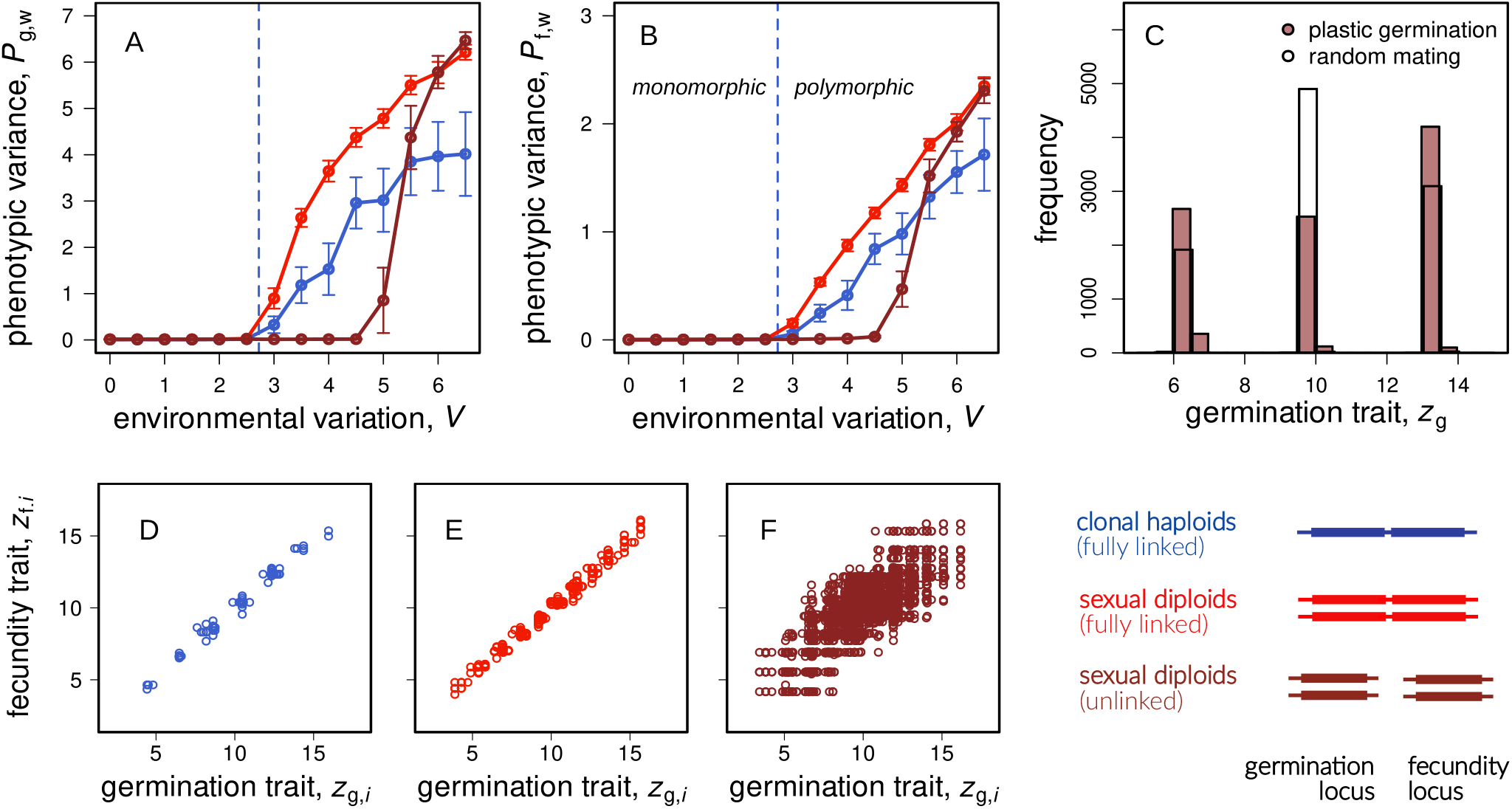
Graphs **A-F** show simulation results for the *two-trait case* with clonally reproducing haploid plants (in blue) and for outcrossing diploid plants with and without genetic linkage (light and dark red curves, respectively). The vertical dashed lines in graphs A and B present the analytical threshold between conditions leading to monomorphic and polymorphic plant populations (from the Hessian matrix C6 with components C17-C19). The whiskers represent variation among replicates (spanning one standard deviation) and the parameter values for these simulations are: 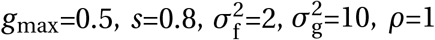. The histograms in graph **C** present the phenotypic distribution at the end of a single simulation run with temporal assortative mating due to germination plasticity (dark red bars) and with random mating among all plant seeds (transparent bars). Temporal assortative mating leads to a reduced hybridization between the outer morphs relative to the case of random mating. Simulation parameters are the same as in graphs A-B with *V* =6. Graphs **D-F** show the phenotypic distribution of single simulation runs for varying genetic architectures with parameter values: 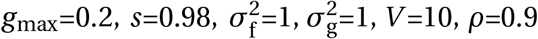.

We further investigated how closely the phenotypic distributions of germinated plants matched the underlying environmental distributions (Fig. 1B). First, we measured the phenotypic covariance (*P*_gf,w_; see eq. D7) between individual germination and fecundity traits (*z*_g,*i*_ and *z*_f,*i*_). The sign of this covariance consistently matched the sign of the environmental correlation *ρ* (Fig. 4B), confirming our analytical predictions (eq. C19 in Appendix C). Next, we compared the slope of the linear regression of germination on fecundity traits (*α = P*_gf,w_/*P*_g,w_; Fig. 4C) to the environmental correlation *ρ*. Though *α* and *ρ* were closely aligned, they did not match perfectly (i.e., *α≠ ρ*; Fig. 4D). Such deviations occurred because trait evolution depended not only on adaptation to environmental fluctuations, but also on reducing competition between specialized morphs. As a consequence, plants evolved phenotypic trait combinations that slightly deviated from a perfect match to environmental conditions. Similar results were found by Kortessis and Chesson (2021), who observed reduced predictive germination as a consequence of competition among plants.

We also measured between-year phenotypic variance (*P*_g,b_ and *P*_f,b_; see eq. D6). Polymorphic populations consistently exhibited higher between-year variance (Fig. 4E,F). Moreover, this variance was greater among germinated plants than among seeds in the seed bank (Fig. 4E). This was because polymorphic populations could respond more strongly to changing selective pressures across years, causing rapid changes in morph frequencies that track environmental changes. Under plastic germination, only phenotypes well-adapted to current environmental conditions germinated each year, even when the composition of the seed bank remained relatively stable. Thus, the phenotypic composition of germinated plants became uncoupled from that of the seed bank, leading to substantial year-to-year shifts. Between-year variance among germinated plants accounted for up to approximately 30% of the total phenotypic variance (Fig. 4E), indicating that a single-year measurement captured only about 70% of the total long-term phenotypic variance.

### 3.4 Outcrossing and the rise of temporal assortative mating

As a final extension, we examined the evolution of polymorphism in diploid plants with sexual reproduction. Here, germination and fecundity traits were controlled by separate genetic loci without physical linkage or pleiotropy, and mating was random. Under these conditions, polymorphism evolved under narrower environmental conditions compared to clonal reproduction (compare blue and dark red lines, Fig. 5A,B). This was because recombination breaks down genetic associations between germination and fecundity traits, which were necessary for adaptation to correlated environments (*ρ≠* 0). Consistent with this interpretation, when loci controlling germination and fecundity were completely linked, polymorphism evolved under conditions similar to those with clonal reproduction (light red lines, Fig. 5A,B).

However, when environmental fluctuations were sufficiently strong, polymorphism emerged even without physical linkage between loci. This occurred through the evolution of temporal assortative mating driven by divergence in germination timing (*z*_g_, dark red curves in Fig. 5A,B). Plants with differentiated germination traits germinated preferentially in different years, thus reducing gene flow between differentiated morphs. This temporal segregation enabled genetic associations (linkage disequilibrium) between germination and fecundity traits to establish despite free recombination (Fig. 5F). Temporal assortative mating was also reflected clearly in the phenotypic distribution of the population (Fig. 5C). Under sexual reproduction, two highly differentiated morphs coexisted, along with an intermediate morph that resulted from mating between them. However, this intermediate morph occurred less frequently than would be expected if matings were entirely random (Fig. 5C, red vs. white histograms).

Phenotypic variance was typically larger in polymorphic populations under sexual reproduction compared to clonal reproduction, in particular in the germination trait (Fig. 5A, dark red curves exceed blue curves for large *V*). This was because divergence in the germination trait (*z*_g_) was under additional selection in sexual populations as differentiated germination schedules led to temporal assortative mating, reducing the formation of maladapted hybrids (see Aubier et al., 2023, for a similar result). Additionally, sexual reproduction further led to a so-called segregation variance (Slatkin and Lande, 1994) which also contributed to greater *P*_g,w_ and *P*_f,w_ in our simulations with sexual reproduction (compare blue to red curves in Fig. 5A and B).

## 4 Discussion

Our analyses indicate that adaptive plasticity in seed germination readily evolves and facilitates the emergence and maintenance of genetic diversity in annual plants under temporally fluctuating environments. Such predictive germination boosts the storage effect by allowing seeds to germinate preferentially in years favorable for growth and reproduction. This in turn promotes divergence among genetic morphs that can specialize to different yearly conditions and leads to temporal niche partitioning. In contrast, when genetic constraints lead to maladaptive plasticity, seeds often germinate into unfavorable conditions, opposing diversification. Our results thus extend existing theory on the storage effect (Venable et al., 1993; Snyder and Adler, 2011; Kortessis and Chesson, 2021), demonstrating that adaptive plasticity and genetic polymorphism can interact to reinforce biodiversity.

Predictive germination facilitates diversity by generating positive correlations between germination timing and favorable conditions for reproduction. When seeds reliably detect cues predicting good years, they can align their germination decisions with environments best suited for growth and fecundity. Such alignment reduces competition among emerging genetic morphs, as each specializes on a distinct subset of years (Chesson and Warner, 1981; Ellner and Hairston, 1994). This interplay between germination and fecundity traits produces strong phenotypic covariance, further promoting niche partitioning and enabling stable coexistence (Kortessis and Chesson, 2021). Adaptive germination plasticity thus enhances temporal niche differentiation beyond what is possible with divergence in a single trait.

Our results also suggest that predictive germination can promote reproductive isolation, thus setting the stage for ecological speciation. Divergence in germination timing generates temporal assortative mating, reducing gene flow among plant morphs specialized on different annual conditions. In that sense, the germination trait in our model can be seen as a “magic trait”, simultaneously controlling adaptation to the environment and facilitating reproductive isolation (Servedio et al., 2011). Such traits are particularly effective drivers of ecological speciation because they prevent genetic recombination and sexual reproduction from breaking down adaptive trait combinations (Dieckmann and Doebeli, 1999; Gavrilets and Vose, 2007). Even when loci controlling germination and fecundity traits are unlinked, temporal assortative mating allows trait covariance to emerge and persist, circumventing recombination’s homogenizing effects (Dieckmann and Doebeli, 1999; Aubier et al., 2023). When temporal assortative mating via plastic germination is insufficient to override recombination, selection may favor tighter genetic linkage or pleiotropy between germination and post-germination traits, potentially promoting “supergenes” that further stabilize adaptive trait combinations (Sinervo and Svensson, 2002). Alternatively, other mechanisms strengthening temporal assortative mating, such as genetic incompatibility systems, could enhance reproductive isolation, avoid the production of unfit hybrids, and thus facilitate diversification.

Our findings contribute to a broader discussion on how phenotypic plasticity affects genetic diversification. Adaptive plasticity is often viewed as an alternative adaptation to genetic diversification, potentially limiting genetic divergence by enabling a single genotype to exploit multiple environments (Ghalambor et al., 2007; Pfennig et al., 2010; Turcotte and Levine, 2016; Schmid and Guillaume, 2017). Predictive germination provides an interesting exception, showing instead how plasticity can facilitate genetic diversification. This illustrates a more general mechanism whereby the existence of traits or traits’ associations that allow genotypes to be preferentially expressed in contexts where they perform best favors diversification (Avila and Mullon, 2023; Lehmann and Mullon, 2025). Our arguments therefore also apply more broadly to life-history plasticity, where adaptive transitions between life stages may allow populations to exploit distinct temporal niches more effectively than divergence in a single trait alone. Rather than being mutually exclusive, adaptive plasticity and genetic diversification can thus act synergistically to enhance biodiversity (see also West-Eberhard 1989 for an alternative mechanism; Hendry 2016 and Sommer 2020 for a review).

Empirical studies provide support for our findings on predictive germination. For example, some long-term studies of desert annual plants indicate associations between germination timing and conditions favorable for reproduction, such as specific rainfall or temperature cues (Venable et al., 1993; Pake and Venable, 1996; Gremer et al., 2016). Also, germination fractions within a year seem to correlate with the expected competitive environment (Tielbörger and Valleriani, 2005; Tielbörger and Prasse, 2009). Other potentially relevant examples come from fire-prone ecosystems, where germination cues such as smoke or heat may reliably predict beneficial growth conditions, including enhanced nutrient availability and reduced competition (Keeley and Pausas, 2022). In these systems, germination traits correlate with adaptive post-germination traits like high-light tolerance or rapid post-fire growth (Pausas and Lamont, 2022).

Our findings also provide a potential explanation for the pronounced between-year variation observed in some annual plant communities. In our model, predictive germination creates strong year-to-year changes in the phenotypic composition of germinated plants, despite a relatively stable composition of the underlying seed bank. Such between-year fluctuations in community composition have indeed been reported from long-term field surveys, with some species appearing abundantly in certain years and remaining virtually undetected in others (Pake and Venable, 1996; Facelli et al., 2005; Levine et al., 2008; Tielbörger et al., 2014; Carrasco-Puga et al., 2021). These dynamics could complicate biodiversity assessments because short-term surveys may underestimate true diversity hidden in dormant seed banks (Carrasco-Puga et al., 2021). Accurate evaluation of biodiversity under temporal fluctuations therefore calls either for long-term monitoring data, comprehensive germination trials, or genetic techniques such as environmental DNA analyses to fully capture community diversity.

Climate change could alter the ecological and evolutionary implications of predictive germination by modifying patterns of environmental predictability. As environmental cues may become less reliable with climate change (Kingsolver and Buckley, 2018; Visser and Gienapp, 2019; Edwards and Yang, 2021), the capacity of seeds to match germination timing to optimal growth conditions may decline, reducing the effectiveness of temporal niche partitioning and potentially eroding biodiversity. In particular, increasingly erratic rainfall patterns, temperature shifts, and altered disturbance regimes could disrupt previously stable associations between germination cues and favorable reproductive conditions. Such disruptions could disproportionately affect plant communities adapted to strongly correlated environmental fluctuations, possibly leading to declines in population sizes and shifts in species composition. Understanding how predictive germination strategies might respond to ongoing environmental change therefore emerges as a relevant question for future biodiversity dynamics.

Because our model is based on several simplifying assumptions, further modifications might be needed to tailor the model to specific systems. For instance, seed densities in the soil can vary greatly between years (while we assume constant *N*) and germination fractions in desert annuals might predominantly increase with precipitation (e.g., see Lampei et al., 2017, while we assume a bell-shaped reaction norm). In addition, the model does not consider parental effects (Tielbörger and Petrů, 2010) even though this is known to contribute to temporal niche partitioning (Yamamichi and Hoso, 2017). Beside these ecological aspects, seed germination and adult fecundity often posses polygenic trait architectures (while we modeled a one-locus-one-trait architecture), that might further complicate the buildup of trait covariances (e.g., Chebib and Guillaume, 2022). Extending our model for such alternative ecological and genetic properties might modify the evolutionary outcome and thus could offer additional insights into the evolution of temporal niche partitioning in annual plants.

To conclude, our study reveals that adaptive plasticity in seed germination promotes genetic diversification by facilitating temporal niche partitioning and reproductive isolation in temporally fluctuat-ing environments. These results highlight that adaptive plasticity and genetic diversification are not mutually exclusive but instead can act together to enhance biodiversity. Understanding such interactions will help us better understand how natural populations might respond to future environmental change.

## 5 Acknowledgement

A.D. was funded by the SAGE Centre, a Global Climate Centre initiative of the German Academic Exchange Service (DAAD).

### A Appendix - Environmental variation

#### A.1 Bivariate normal distribution

The environmental conditions during germination and reproduction (***θ***_*t*_ *=* (*θ*_g,*t*_, *θ*_f,*t*_)) for each year *t* are picked from a Bivariate normal distribution

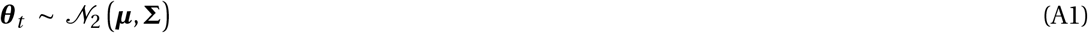

with the vector of average environmental conditions

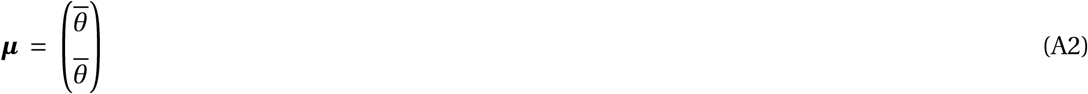

and environmental variance-covariance matrix

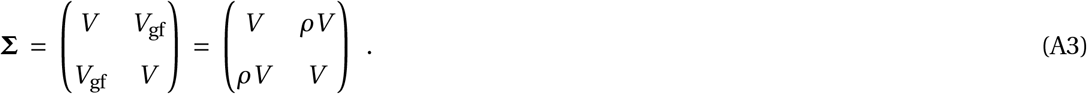

The environmental distribution thus is characterized by between-year environmental variance *V* (see eq. A7), the covariance between the germination and fecundity environment *V*_gf_ (see eq. A8), and the within-year correlation coefficient *ρ*.

With a bivariate normal distribution, the probability density function of *θ*_g,*t*_ and *θ*_f,*t*_ is

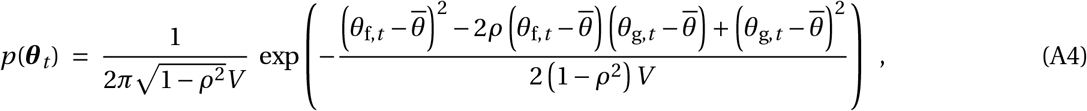

where *θ*_g,*t*_ and *θ*_f,*t*_ are correlated to each other with correlation coefficient *ρ*. When considering the long-term distribution of each environmental value separately, both *θ*_g,*t*_ and *θ*_f,*t*_ follow a univariate normal distribution with probability density function

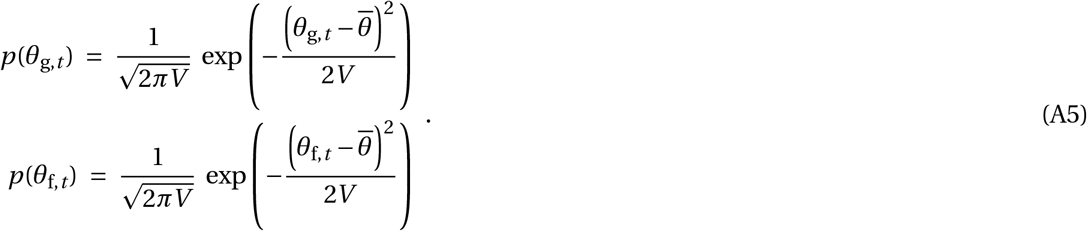

Both environmental conditions share the same average value *θ* across years

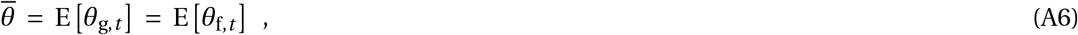

and the same between-year variance *V*

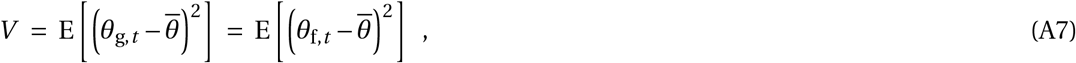

where expectations are taken over *p*(***θ***_*t*_) throughout. The covariance among the germination environment *θ*_g,*t*_ and the reproduction environment *θ*_f,*t*_ is

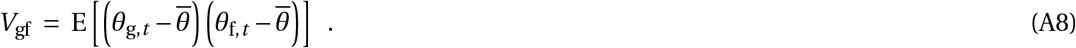

#### A.2 Conditional distribution

To illustrate how environmental conditions change within each year, we provide the conditional distribution of the fecundity environment *θ*_f,*t*_ given a certain value of the germination environment *θ*_g,*t*_ and within-year environmental correlation *ρ*. This conditional distribution follows the univariate normal distribution

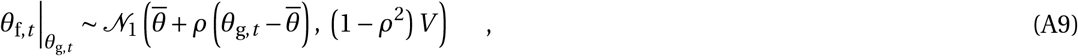

illustrating that the degree of environmental correlation *ρ* re-scales the conditional mean 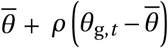 and the conditional variance 1 *−ρ*^2^ *V*. In the extreme case of *ρ =* 1, the conditional distribution becomes

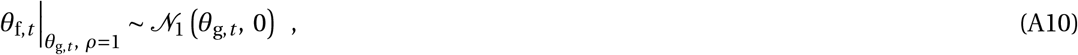

and in the case of *ρ = −*1 reduces to

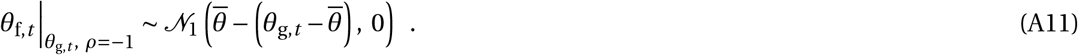

These two cases illustrate that a large positive environmental correlation leads to constant environmental conditions within years (i.e. *θ*_g,*t*_ *≈ θ*_f,*t*_ with *ρ →* 1), while large negative environmental correlation installs large within-year environmental changes (i.e. 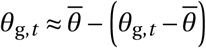 with *ρ → −*1).

### B. Appendix - Single-trait case

#### B.1 Analysis

We first investigate the *single-trait case*, where germination and fecundity are controlled by the same trait (*z*_g,*i*_ *= z*_f,*i*_ *= z*_*i*_). We study the evolution of this trait using an evolutionary invasion analysis. To this end, we consider a rare mutant with trait value *z*_m_ appearing in a large population that is otherwise monomorphic for a resident trait *z* (with mutational effect being small such that *z*_m_ *−z* is small). Whether or not the mutant can invade and substitute the resident trait value, can be determined from the mutants’ invasion fitness (i.e. its geometric growth rate). Here we’ll use the invasion fitness proxy (i.e. sign equivalent around one)

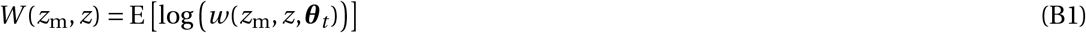

(e.g., see Lewontin and Cohen, 1969; Sæther and Engen, 2015) where

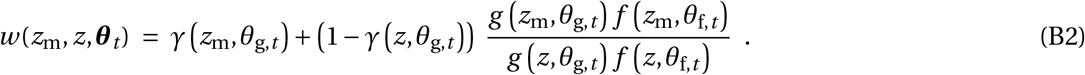

is individual fitness, which here is the expected number of seeds that a mutant seed in the seed bank at the beginning of year *t* contributes to next year’s seed bank. Individual fitness (eq. B2) is composed of two parts reflecting two potential contributions. First, the mutant seed may stay in the seed bank (i.e. not germinate) and survive there to the next year, which occurs with probability

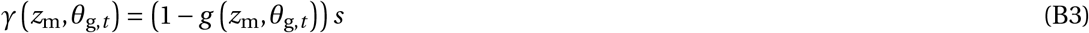

(where *g z*_m_, *θ*_g,*t*_ is given by eq. 1 in the main text). The second term of eq. (B2), meanwhile, captures that the mutant seed may germinate, reproduce with fecundity *f z*_m_, *θ*_f,*t*_ (given by eq. 2 in the main text), and contribute new seeds to the seed bank of the next year, competing with residents for 1 *−γ z, θ*_g,*t*_ sites. See Chesson and Warner (1981) and Svardal et al. (2011) for similar formulations of individual fitness.

The function *W* (*z*_m_, *z*) can be used to infer on long term evolution via a sequence of invasions of mutations. The selection gradient

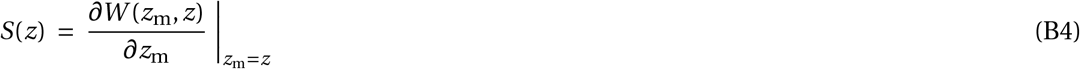

indicates directional selection during this sequence, with selection favoring trait increase when *S*(*z*) *>* 0 and trait decrease when *S*(*z*) *<* 0. A strategy *z*^*∗*^ where *S*(*z*^*∗*^) *=* 0 is called a singular strategy. Whether or not such a singular strategy will be approached by gradual evolution is determined by the sign of

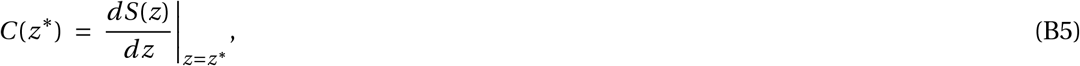

with *C* (*z*^*∗*^) *<* 0 indicating that *z*^*∗*^ is an attractor (i.e. is convergence stable) and *C* (*z*^*∗*^) *>* 0 indicating that *z*^*∗*^ is a repellor of evolutionary dynamics. Once a population has converged to an attractor *z*^*∗*^, the disruptive selection coefficient

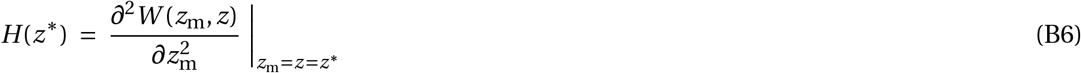

determines whether the population remains monomorphic (when *H* (*z*^*∗*^) *<* 0) or becomes polymorphic in a process referred to as evolutionary branching (when *H* (*z*^*∗*^) *>* 0).

#### B.2 Results

##### B.2.1 Selection gradient

Substituting eq. (B1) into eq. (B4), we obtain the selection gradient

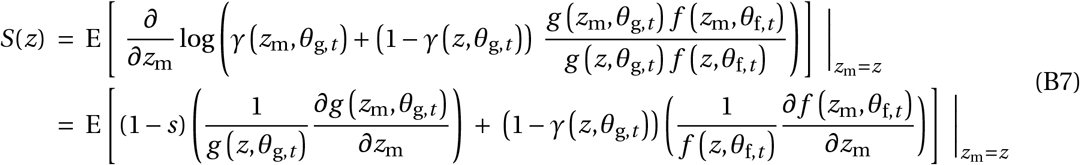

with the derivatives being

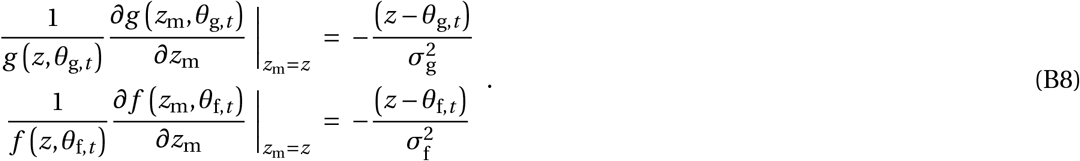

To average over years, we recall that 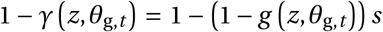 and compute the long-term average germination probability 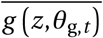 as

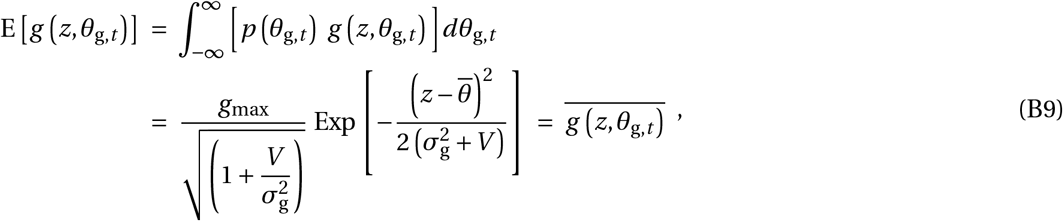

and the average product of germination probability and the environment during adult fecundity 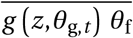 as

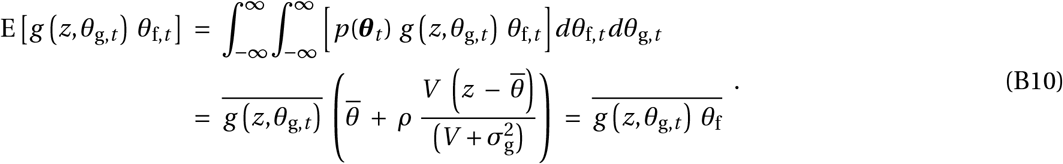

When plugging eqs. (B8)-(B10) into eq. (B7), the selection gradient becomes

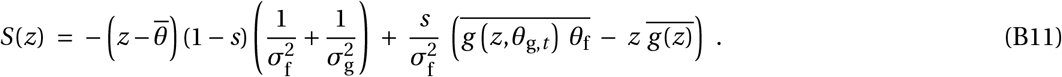

Solving this for zero shows that the only singular strategy is 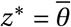.This singular strategy is convergence stable because

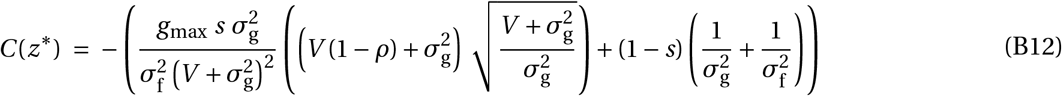

is always negative.

##### B.2.2 Disruptive selection coefficient

We plug the expression for invasion fitness (eq. B1) into the expression for the disruptive selection coefficient (eq. B6) and obtain

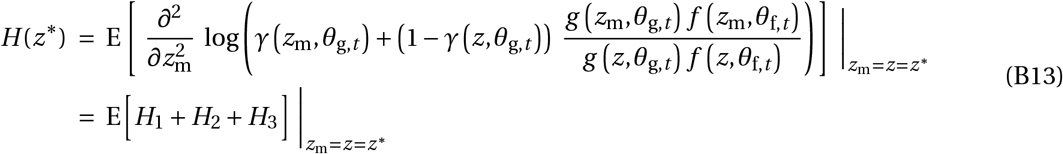

with components

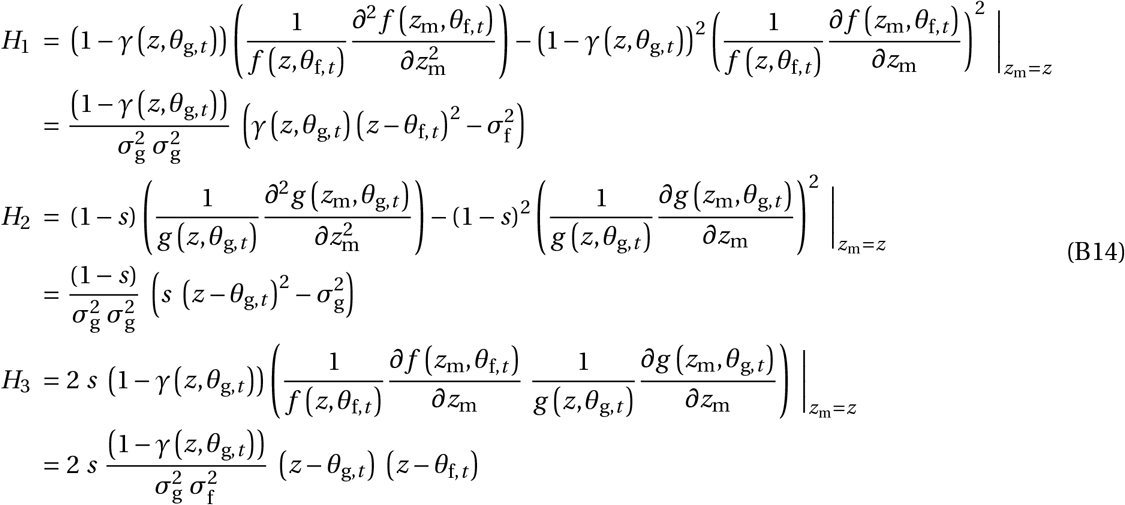

To average over infinitely many years, we compute the long-term averages at the singular strategy as

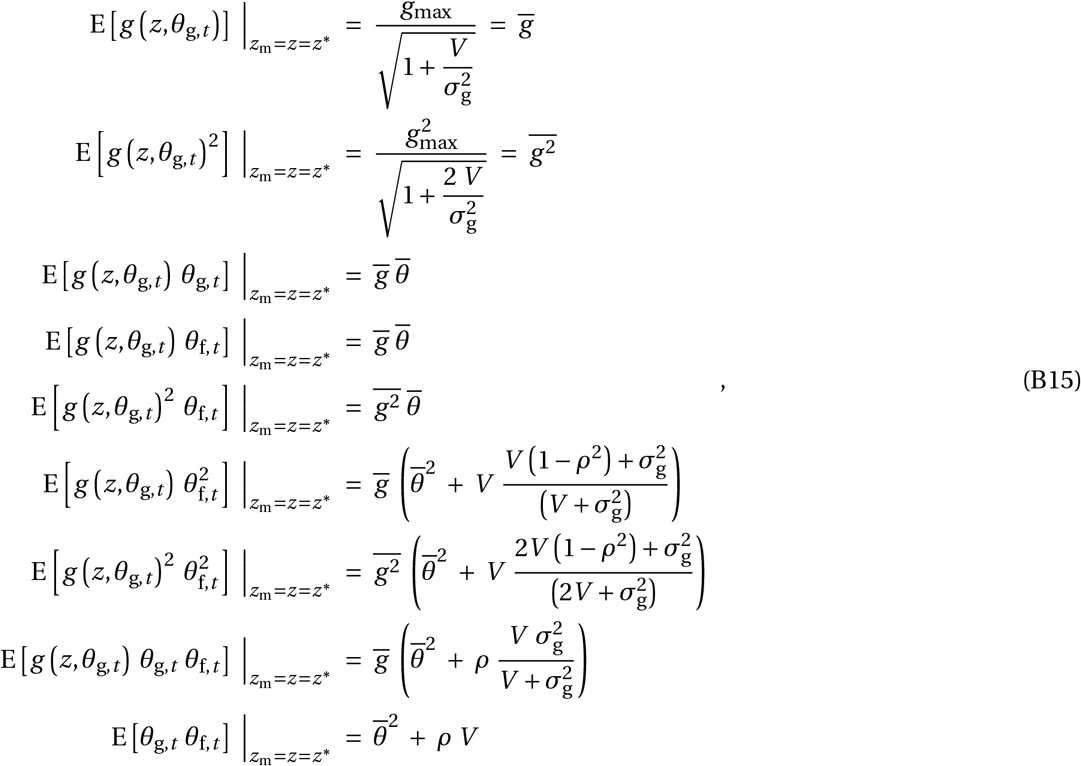

where expectations in eq. (B15) are taken over *p*(***θ***_*t*_), similar to the approach in eq. (B10). We then compute the components of *H* (*z*^*∗*^) as

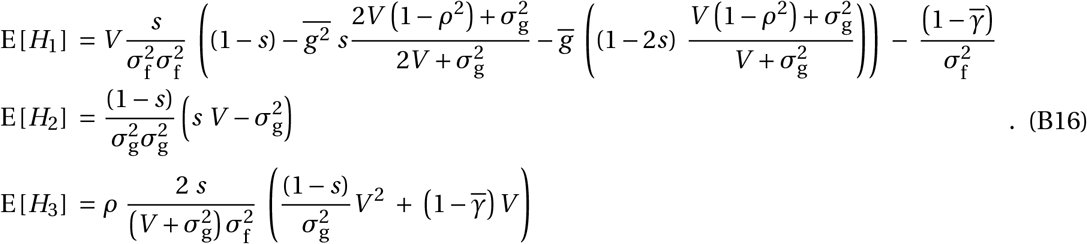

Expressions (B16) show that greater germination plasticity (i.e. smaller 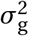), in combination with selection on adult fecundity (i.e. with finite 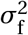), can either facilitate or oppose trait polymorphism (i.e. either increase or decrease *H* (*z*^*∗*^)). On the one hand, greater germination plasticity reduces the negative terms in components E[*H*_1_] and E[*H*_2_], and thus increases the overall disruptive selection coefficient *H* (*z*^*∗*^). Specifically, a smaller 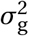 increases E[*H*_1_] because germination plasticity reduces the long-term average germination probabilities 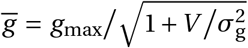 and 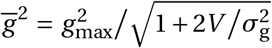 (see eqs. B15) and thus increases the generation overlap 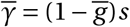. On the other hand, greater germination plasticity increases the magnitude of component E[*H*_3_] (when a smaller 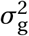 in the denominator within the bracket increases E[*H*_3_]), where the sign of E[*H*_3_] depends on the sign of environmental correlation *ρ*. Germination plasticity thus leads to a larger negative component E[*H*_3_] with negative environmental correlation *ρ* (and thus strengthens stabilizing selection), but leads to a larger positive component E[*H*_3_] with positive environmental correlation *ρ* (and thus strengthens disruptive selection). This behavior explains mathematically how germination plasticity promotes trait polymorphism with positive *ρ*, but opposes polymorphism with negative *ρ* (i.e. see Fig. 2).

Even though this is not immediately obvious from eq. (B16), the disruptive selection coefficient in the absence of plastic germination 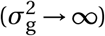 reduces to

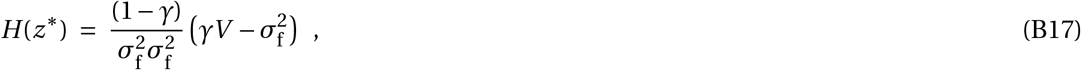

where generation overlap *γ =* (1*−g*_max_)*s*. In the absence of adult selection 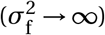, the components E[*H*_1_] and E[*H*_3_] both become zero and

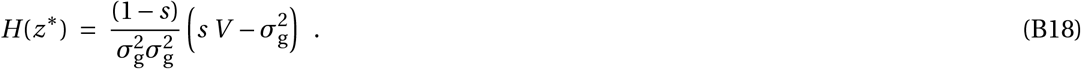

In absence of environmental correlation between germination and reproduction (*ρ =* 0), the disrup-tive selection coefficient reduces to

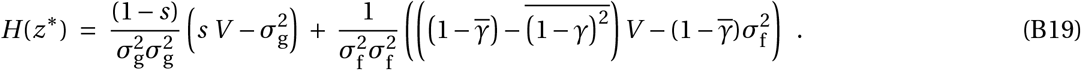

According to eq. (B19), plastic seed germination 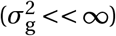 at *ρ =* 0 has a two-fold effect on evolutionary branching when combined with adult selection 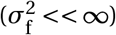. First, plastic germination contributes to the first part of *H* (*z*^*∗*^) (eq. B19) and could either favor or disfavor disruptive selection depending on whether 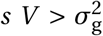 or 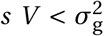. Second, germination plasticity modifies the second part of *H* (*z*^*∗*^) (eq. B19) by reducing generation overlap 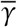 and 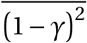. Here, the long-term generation overlap 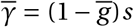 is a function of the long-term germination probability 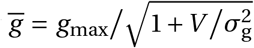 (that is here estimated for a monomorphic population at the singular strategy *z*^*∗*^; see eq. B15) that declines with greater germination plasticity (i.e. with smaller 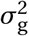). To make a long story short, plasticity in seed germination can either facilitate or oppose niche partitioning at *ρ =* 0 as compared to the non-plastic case (e.g., see Fig. 2 for *ρ =* 0).

### C Appendix - Two-trait case

#### C.1 Analysis

In the *two-trait case*, we closely follow the model of the main text (see section 2) and assume that the germination probability *g* (*z*_g,*i*_, *θ*_g,*t*_) and adult fecundity *f* (*z*_f,*i*_, *θ*_f,*t*_) are controlled by two separate traits, termed *z*_g_ and *z*_f_. These traits are controlled by two distinct loci, and mutations have independent effects on both traits (i.e. mutations only hit one trait at a time). We characterize the resident population by trait vector

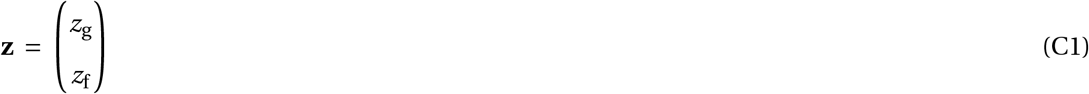

and the mutant individuals by trait vector

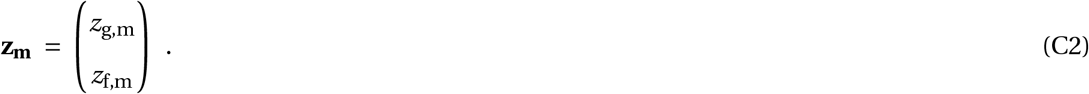

The invasion fitness proxy of a mutant individual in the *two-trait case* (i.e. its geometric growth rate) is

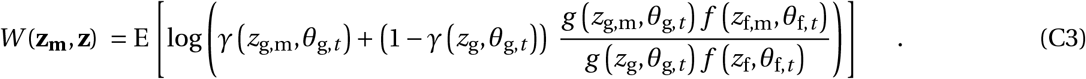

This expression closely follows that of the *single-trait case* (eq. B1 and eq. B2), while two distinct traits control seed germination and fecundity, respectively.

We investigate directional selection on trait *z*_g_ and *z*_f_ by vector

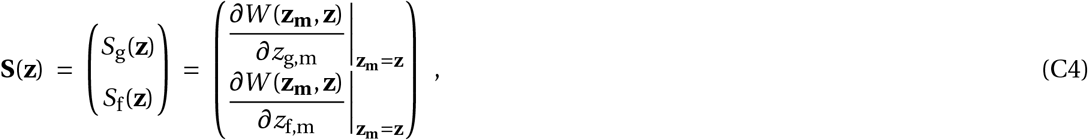

and find directional selection to be absent at the singular strategy 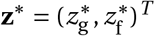 when *S*_g_(**z**^*∗*^) *=* 0 and *S*_f_(**z**^*∗*^) *=* 0. The singular strategy is an attractor of directional trait changes when all eigenvalues of the Jacobian matrix

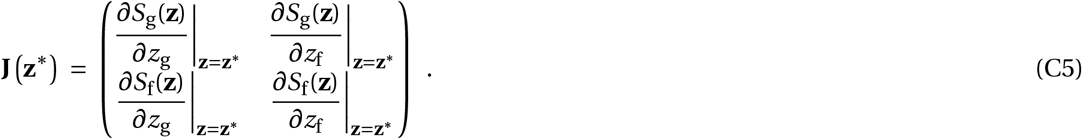

have a negative real part (i.e. **z**^*∗*^ then is convergence stable). Upon approaching **z**^*∗*^, the population becomes polymorphic when the greatest eigenvalue *λ*(**z**^*∗*^) of the Hessian matrix

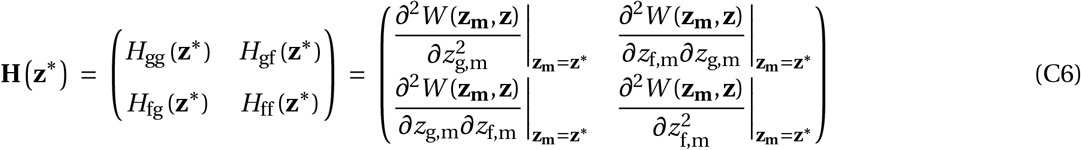

is positive (*λ*(**z**^*∗*^) *>* 0), and the population remains monomorphic at a negative eigenvalue (*λ*(**z**^*∗*^) *<* 0).

#### C.2 Results

##### C.2.1 Selection gradient

Following eqs. (C3) and (C4), the selection gradient in the two-trait model becomes

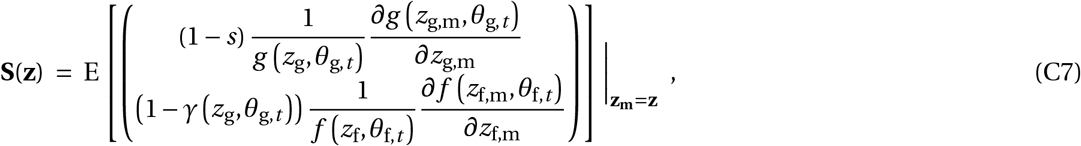

with the derivatives

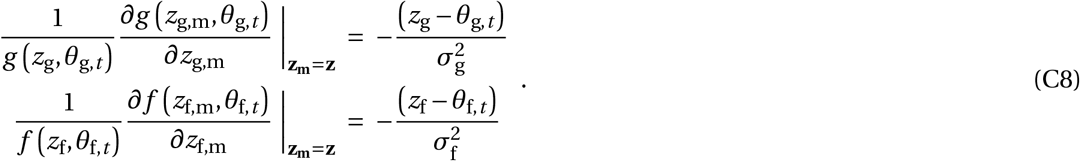

For a population that is monomorphic for the trait vector **z**, the long-term average germination probability 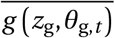 and the long-term average 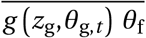 are

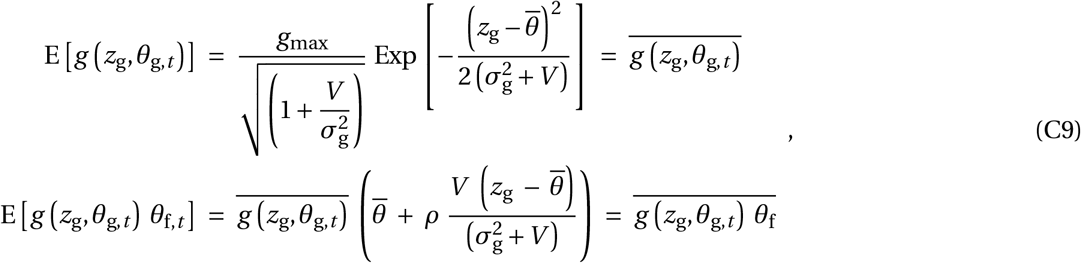

with the expectations being taken over *p*(***θ***_*t*_), respectively. The selection gradient then becomes

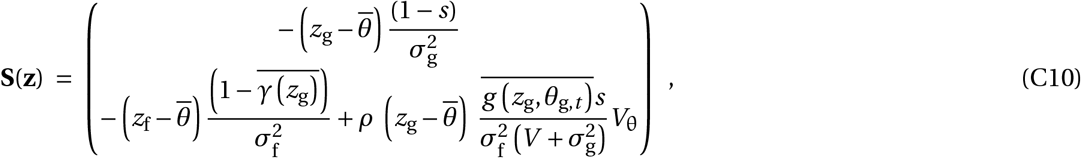

with the singular strategy

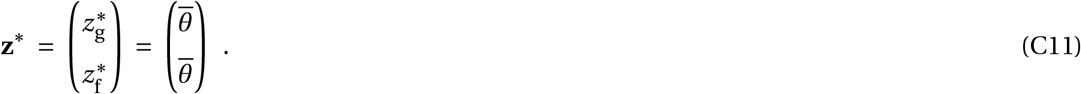

This singular strategy is convergence stable because the eigenvalues of the Jacobian

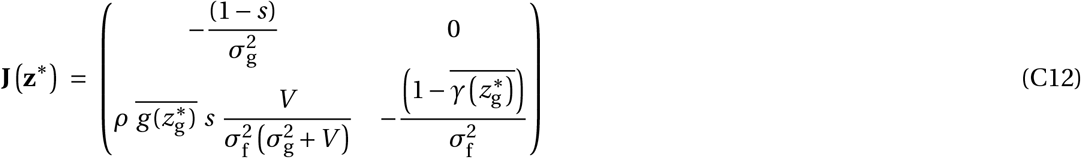

are both negative (i.e. 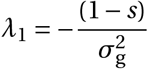 and 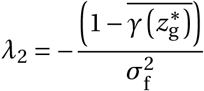).

##### C.2.2 Disruptive selection coefficient

The entries of the Hessian matrix, following expressions (C3) and (C6), become

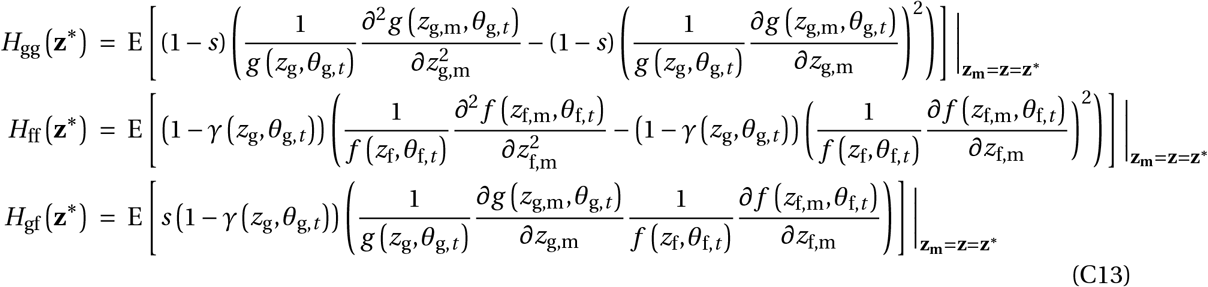

with the first order derivatives following eqs. (C8) and the second order derivatives being

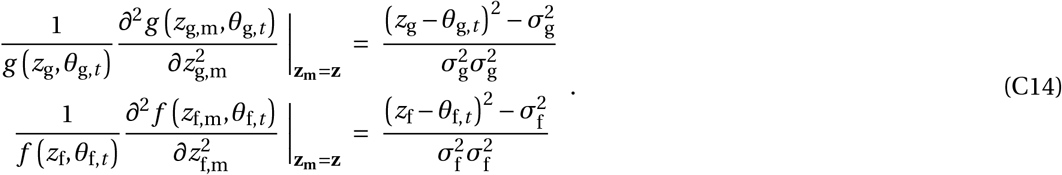

When averaging over infinitely many years for a population that is monomorphic for the singular strategy, we obtain

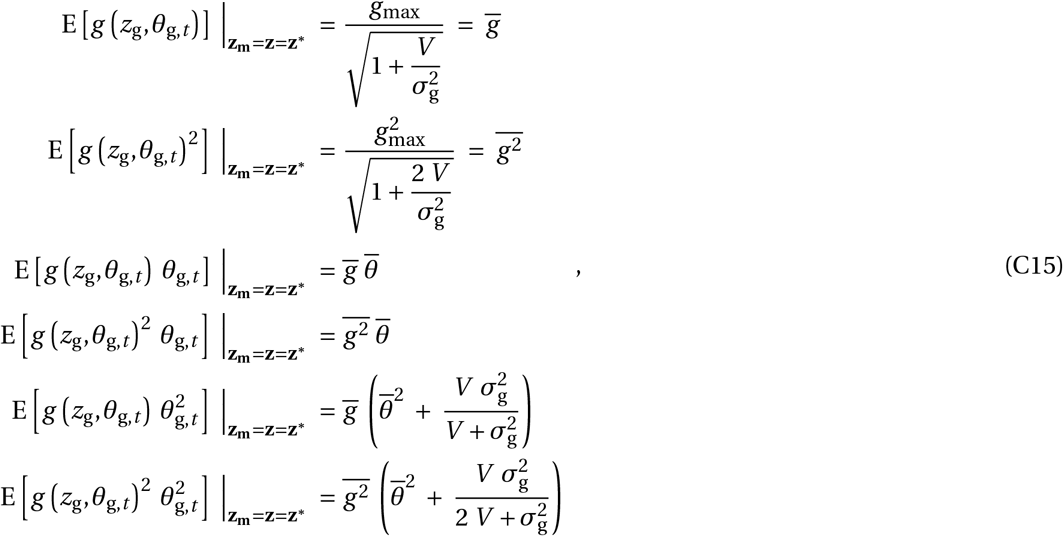

and

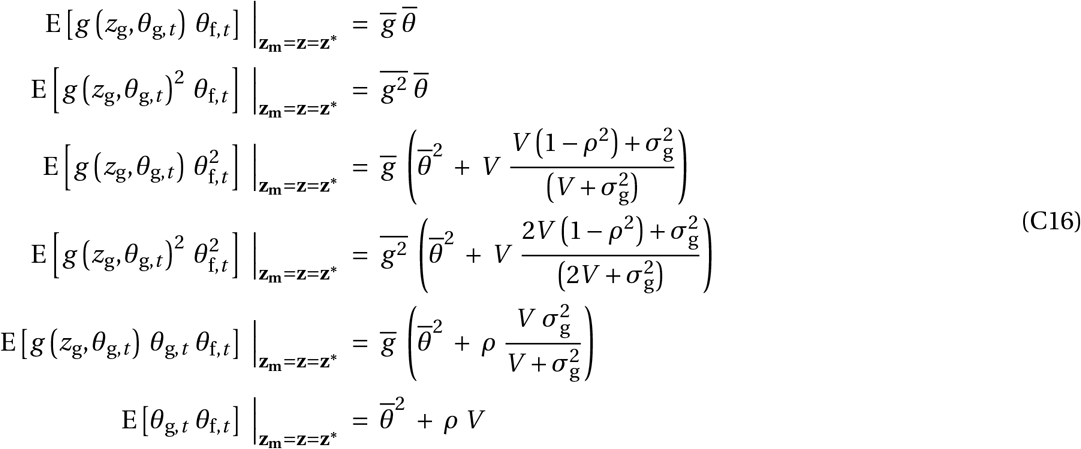

Note that the expectations in eqs. (C15)-(C16) are taken over *p*(***θ***_*t*_), similar to the approach in eq. (B10). The first entry of the Hessian matrix *H*_gg_ (**z**^*∗*^), that is the quadratic coefficient of selection on the germination trait, then becomes

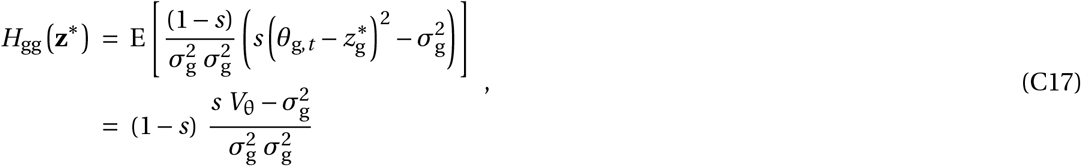

a term that is positive (i.e. *H*_gg_ (**z**^*∗*^) *≥* 0) when 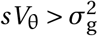. The second diagonal entry of the Hessian, that is the quadratic coefficient of selection on the fecundity trait, is

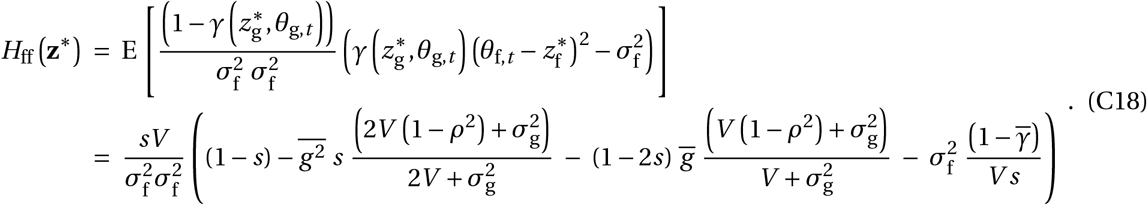

Note that the quadratic selection coefficient on the fecundity trait *H*_ff_ (**z**^*∗*^) is not only a function of fecundity related parameters, like adult niche breadth 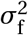, but also changes with germination plastic-ity 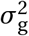. In fact, greater germination plasticity (i.e. smaller 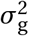) increases coefficient *H*_ff_ because ger-mination plasticity reduces the long-term average germination probability (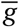 and 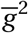, see eqs. B15), and thus promotes generation overlap 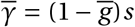. In the absence of germination plasticity (when 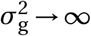), component *H*_ff_ (**z**^*∗*^) reduces to 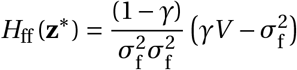 and we recover the classic branch-ing condition 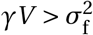 with *γ =* (1*−g*_max_)*s* (e.g., Chesson and Warner, 1981; Ellner and Hairston, 1994; Svardal et al., 2015).

The off-diagonal entry of the Hessian, that is the correlational coefficient of selection, is

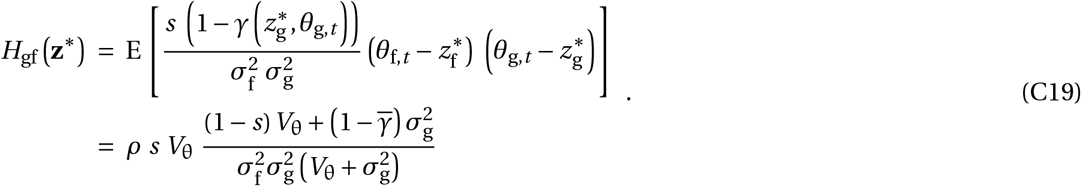

The degree of environmental correlation *ρ* determines the magnitude and sign of correlational selection, and thus determines the kind of trait covariance favored by natural selection. With positive environmental correlation *ρ >* 0, a positive trait covariance evolves because *H*_gf_ (**z**^*∗*^) *>* 0. With a nega-tive environmental correlation (*ρ <* 0), a negative trait covariance between *z*_g_ and *z*_f_ evolves because *H*_gf_ (**z**^*∗*^) *>* 0. More generally, the presence of correlational selection in our model always promotes disruptive selection because

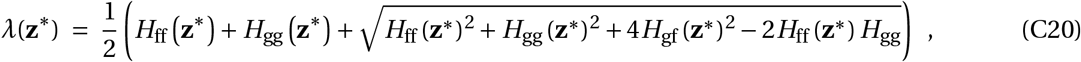

and the absolute value of *H*_gf_ (**z**^*∗*^) always increases *λ*(**z**^*∗*^)).

### D Appendix - Simulations

#### D.1 Genetic basis of traits

To accompany the mathematical analyses of the *two-trait case*, we ran individual-based simulations in *R* for two separate scenarios: 1) clonally reproducing haploid plants, and 2) sexually reproducing diploid plants. The fecundity trait *z*_g,*i*_ and the germination trait *z*_f,*i*_ each was controlled by a separate genetic locus, in the absence of pleiotropy and in absence of any random trait contributions (e.g., there was no developmental noise contributing to *z*_g,*i*_ and *z*_f,*i*_). At the beginning of each simulation, the allelic value at each homologous copy was picked from a normal distribution 𝒩_1_(5, 0.25) for each diploid individual separately (with 𝒩_1_(10, 0.25) in the haploid case). These allelic effects at the homologous copies then were summed up to compute the germination and fecundity trait value *z*_g,*i*_ and *z*_f,*i*_, respectively (i.e. we assumed additive gene actions). Mutations occurred in newly produced seeds with a per-site mutation rate *µ =* 0.00025 in diploids (and a per-locus mutation rate of *µ =* 0.0005 in haploids), at each locus separately. The mutational effects were picked from a normal distribution *𝒩*_1_*(0,0*.*01)* and added to the existing allelic value following the continuum-of-alleles model.

#### D.2 Environmental variation

We picked the environmental conditions during germination and reproduction *θ*_g,*t*_ and *θ*_f,*t*_ each year anew from a bivariate normal distribution

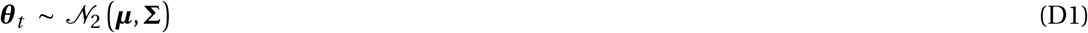

with the vector of average environmental conditions

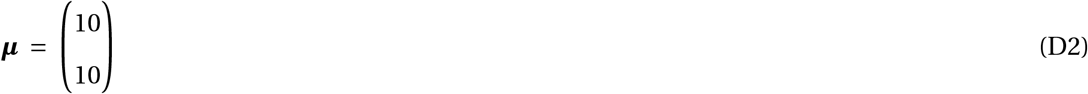

and environmental variance-covariance matrix

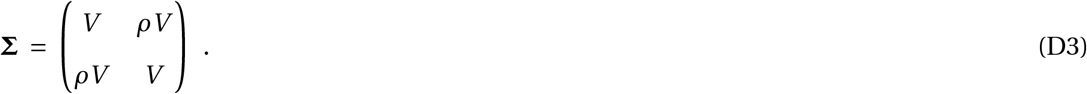

#### D.3 Life-Cycle Events

The life cycle in the simulations matched that one presented in the Model section of the main text. At the beginning of each year, each of the *N* seeds in the seed bank germinated with germination probability *g (z*_g,*i*_, *θ*_g,*t*_) following eq. (1). Each non-germinated seed survived in the seed bank to the next year with probability *s* and died with probability 1 *− s*. Each germinated plant reproduced with individual fecundity *f (z*_f,*i*_, *θ*_f,*t*_) following eq. (2). Newly produced seeds randomly entered the seed bank until *N* seeds were present at the end of each year.

#### D.4 Simulation setting and summary statistics

We ran each simulation for *T* =20 000 years with a total individual number of *N* =10 000 and 20 replicates per parameter combination. At evolutionary equilibrium, we computed the total (long-term) phenotypic variance at the germination and fecundity trait for the last 500 years with a 25-year sample interval (*t =* (19500, 19525, 19550,…, 20000))

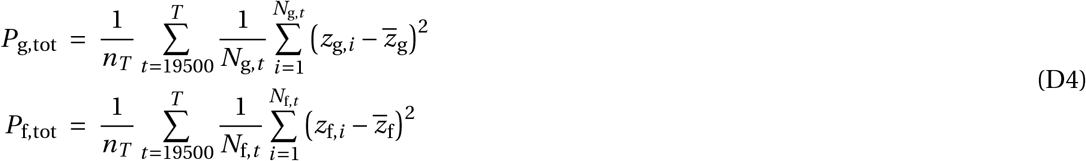

for *n*_*T*_ *=* 21 sampled years, with the long-term average trait value 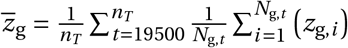 and the number of germinated plants in year *t* being *N*_g,*t*_.

We further partitioned this total phenotypic variance into two variance components, *P*_g,tot_ *= P*_g,w_ *+ P*_g,b_ and *P*_f,tot_ *= P*_f,w_ *+P*_f,b_. First, we computed within-year phenotypic variance

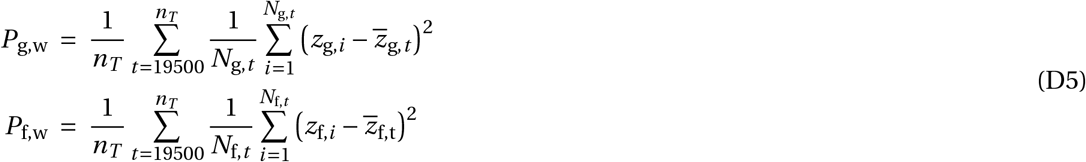

with the average trait value in year *t* being 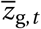. Second, we calculated between-year phenotypic variance

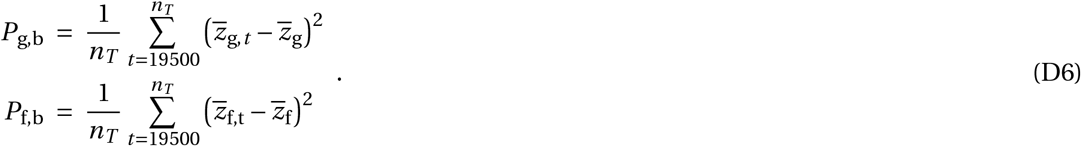

At last, we computed the phenotypic covariance between the germination and fecundity trait within years

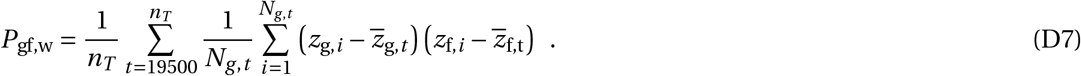

## References

Angert, A. L., Huxman, T. E., Chesson, P., and Venable, D. L. (2009). Functional tradeoffs determine species coexistence via the storage effect. Proceedings of the National Academy of Sciences, 106(28):11641–11645.

Aubier, T. G., Bürger, R., and Servedio, M. R. (2023). The effectiveness of pseudomagic traits in promoting premating isolation. Proceedings of the Royal Society B: Biological Sciences, 290:20222108.

Avila, P. and Mullon, C. (2023). Evolutionary game theory and the adaptive dynamics approach: Adaptation where individuals interact. Philosophical Transactions of the Royal Society B, 378:20210502.

Barabás, G., D’Andrea, R., and Stump, S. M. (2018). Chesson’s coexistence theory. Ecological Monographs, 88(3):277–303.

Bonamour, S., Chevin, L.-M., Charmantier, A., and Teplitsky, C. (2019). Phenotypic plasticity in response to climate change: the importance of cue variation. Philosophical Transactions of the Royal Society B: Biological Sciences, 374(1768):20180178.

Bradshaw, A. D. (1965). Evolutionary significance of phenotypic plasticity in plants. Advances in Genetics, 13:115–155.

Carrasco-Puga, G., Díaz, F. P., Soto, D. C., Hernández-Castro, C., Contreras-López, O., Maldonado, A., Latorre, C., and Gutiérrez, R. A. (2021). Revealing hidden plant diversity in arid environments. Ecography, 44:98–111.

Chebib, J. and Guillaume, F. (2022). The relative impact of evolving pleiotropy and mutational correlation on trait divergence. Genetics, 220(1).

Chesson, P. (2000). Mechanisms of maintenance of species diversity. Annual Review of Ecology and Systematics, 31(1):343–366.

Chesson, P. L. (1983). Coexistence of Competitors in a Stochastic Environment: The Storage Effect. In Levin, S., Freedman, H. I., and Strobeck, C., editors, Population Biology, volume 52, pages 188–198. Springer Berlin Heidelberg, Berlin, Heidelberg. Series Title: Lecture Notes in Biomathematics.

Chesson, P. L. (1994). Multispecies competition in variable environments. Theoretical Population Biology, 45:227–276.

Chesson, P. L. and Warner, R. R. (1981). Environmental variability promotes coexistence in lottery competitive systems. The American Naturalist, 117(6):923–943.

Chevin, L.-M. and Lande, R. (2011). Adaptation to marginal habitats by evolution of increased phenotypic plasticity. Journal of Evolutionary Biology, 24(7):1462–1476.

Cohen, D. (1967). Optimizing reproduction in a randomly varying environment when a correlation may exist between the conditions at the time a choice has to be made and the subsequent outcome. Journal of Theoretical Biology, 16(1):1–14.

Conover, D. O., Duffy, T. A., and Hice, L. A. (2009). The covariance between genetic and environmental influences across ecological gradients: Reassessing the evolutionary significance of countergradient and cogradient variation. Annals of the New York Academy of Sciences, 1168:100–129.

Crispo, E. (2008). Modifying effects of phenotypic plasticity on interactions among natural selection, adaptation and gene flow. Journal of Evolutionary Biology, 21(6):1460–1469.

Day, T. (2000). Competition and the effect of spatial resource heterogeneity on evolutionary diversification. The American Naturalist, 155(6):790–803.

DeWitt, T. J., Sih, A., and Wilson, D. S. (1998). Costs and limits of phenotypic plasticity. Trends in Ecology and Evolution, 13(2):77–81.

Dieckmann, U. and Doebeli, M. (1999). On the origin of species by sympatric speciation. Nature, 400(6742):354–357.

Donohue, K., Rubio De Casas, R., Burghardt, L., Kovach, K., and Willis, C. G. (2010). Germination, postgermination adaptation, and species ecological ranges. Annual Review of Ecology, Evolution, and Systematics, 41(1):293–319.

Dupont, L., Thierry, M., Zinger, L., Legrand, D., and Jacob, S. (2024). Beyond reaction norms: the temporal dynamics of phenotypic plasticity. Trends in Ecology & Evolution, 39(1):41–51.

Edwards, C. B. and Yang, L. H. (2021). Evolved phenological cueing strategies show variable responses to climate change. The American Naturalist, 197(1):E1–E–16.

Ellner, S. P. and Hairston, N. G. (1994). Role of overlapping generations in maintaining genetic variation in a fluctuating environment. The American Naturalist, 143(3):403–417.

Facelli, J. M., Chesson, P., and Barnes, N. (2005). Differences in seed biology of annual plants in arid lands: A key ingredient of the storage effect. Ecology, 86(11):2998–3006.

Gavrilets, S. and Vose, A. (2007). Case studies and mathematical models of ecological speciation. 2. Palms on an oceanic island. Molecular Ecology, 16(14):2910–2921.

Geritz, S. A. H., Kisdi, E., Meszéna, G., and Metz, J. A. J. (1998). Evolutionary singular strategies and the adaptive growth and branching of the evolutionary tree. Evolutionary Ecology, 12:35–57.

Ghalambor, C. K., Hoke, K. L., Ruell, E. W., Fischer, E. K., Reznick, D. N., and Hughes, K. A. (2015). Non-adaptive plasticity potentiates rapid adaptive evolution of gene expression in nature. Nature, 525:372–375.

Ghalambor, C. K., McKay, J. K., Carroll, S. P., and Reznick, D. N. (2007). Adaptive versus non-adaptive phenotypic plasticity and the potential for contemporary adaptation in new environments. Functional Ecology, 21:394–407.

Gotthard, K. and Nylin, S. (1995). Adaptive plasticity and plasticity as an adaptation: A selective review of plasticity in animal morphology and life history. Oikos, 74(1):3.

Gremer, J. R., Kimball, S., and Venable, D. L. (2016). Within-and among-year germination in Sonoran Desert winter annuals: Bet hedging and predictive germination in a variable environment. Ecology Letters, 19(10):1209–1218.

He, T., Lamont, B. B., and Pausas, J. G. (2019). Fire as a key driver of Earth’s biodiversity. Biological Reviews, 94(6):1983–2010.

Hendry, A. P. (2016). Key questions on the role of phenotypic plasticity in eco-evolutionary dynamics. Journal of Heredity, 107:25–41.

Johnson, E. C. and Hastings, A. (2022). Towards a heuristic understanding of the storage effect. Ecology Letters, 25(11):2347–2358.

Keeley, J. E. and Pausas, J. G. (2022). Evolutionary ecology of fire. Annual Review of Ecology, Evolution, and Systematics, 53(1):203–225.

Kingsolver, J. G. and Buckley, L. B. (2018). How do phenology, plasticity and evolution determine the fitness consequences of climate change for montane butterflies? Evolutionary Applications, 11:1231–1244.

Kisdi, E. (2002). Dispersal: Risk spreading versus local adaptation. The American Naturalist, 159(6):579–596.

Kortessis, N. and Chesson, P. (2021). Character displacement in the presence of multiple trait differences: Evolution of the storage effect in germination and growth. Theoretical Population Biology, 140:54–66.

Lampei, C., Metz, J., and Tielbörger, K. (2017). Clinal population divergence in an adaptive parental environmental effect that adjusts seed banking. New Phytologist, 214:1230–1244.

Lande, R. (2014). Evolution of phenotypic plasticity and environmental tolerance of a labile quantitative character in a fluctuating environment. Journal of Evolutionary Biology, 27(5):866–875.

Lehmann, L. and Mullon, C. (2025). Evolution of quantitative traits: exploring the ecological, social and genetic bases of adaptive polymorphism. bioRxiv, pages 2025–06.

Levine, J. M., McEachern, K. A., and Cowan, C. (2008). Rainfall effects on rare annual plants. Journal of Ecology, 96(4):795–806.

Lewontin, R. C. and Cohen, D. (1969). On population growth in a randomly varying environment. Proceedings of the National Academy of Sciences, 62(4):1056–1060.

Miller, E. T. and Klausmeier, C. A. (2017). Evolutionary stability of coexistence due to the storage effect in a two-season model. Theoretical Ecology, 10(1):91–103.

Moran, N. A. (1992). The evolutionary maintenance of alternative phenotypes. The American Naturalist, 139(5):971–989.

Ohtsuki, H., Rueffler, C., Wakano, J. Y., Parvinen, K., and Lehmann, L. (2020). The components of directional and disruptive selection in heterogeneous group-structured populations. Journal of Theoretical Biology, 507:110449.

Orive, M. E., Barfield, M., and Holt, R. D. (2023). Partial clonality expands the opportunity for spatial adaptation. The American Naturalist, 202(5):681–698.

Pake, C. E. and Venable, D. L. (1996). Seed banks in desert annuals: Implications for persistence and coexistence in variable environments. Ecology, 77(5):1427–1435.

Pausas, J. G. and Lamont, B. B. (2022). Fire-released seed dormancy - a global synthesis. Biological Reviews, 97(4):1612–1639.

Pfennig, D. W., Wund, M. A., Snell-Rood, E. C., Cruickshank, T., Schlichting, C. D., and Moczek, A. P. (2010). Phenotypic plasticity’s impacts on diversification and speciation. Trends in Ecology and Evolution, 25(8):459–467.

Saltini, M., Vasconcelos, P., and Rueffler, C. (2023). Complex life cycles drive community assembly through immigration and adaptive diversification. Ecology Letters, 26:1084–1094.

Scheiner, S. M. (1993). Genetics and evolution of phenotypic plasticity. Annual Review of Ecology and Systematics, 24:35–68.

Schlichting, C. D. (1986). The evolution of phenotypic plasticity in plants. Annual Review of Ecology and Systematics, 17(1):667–693. ISBN: 0066-4162.

Schmid, M. and Guillaume, F. (2017). The role of phenotypic plasticity on population differentiation. Heredity, 119(4):214–225. Publisher: Nature Publishing Group.

Servedio, M. R., Doorn, G. S. V., Kopp, M., Frame, A. M., and Nosil, P. (2011). Magic traits in speciation: ‘magic’ but not rare? Trends in Ecology & Evolution, 26(8):389–397.

Simons, A. M. (2014). Playing smart vs. playing safe: The joint expression of phenotypic plasticity and potential bet hedging across and within thermal environments. Journal of Evolutionary Biology, 27(6):1047–1056.

Sinervo, B. and Svensson, E. (2002). Correlational selection and the evolution of genomic architecture. Heredity, 89(5):329–338.

Slatkin, M. and Lande, R. (1994). Segregation variance after hybridization of isolated populations. Genetical Research, 64(1):51–56.

Snyder, R. E. and Adler, P. B. (2011). Coexistence and coevolution in fluctuating environments: Can the storage effect evolve? The American Naturalist, 178(4):E76–E84.

Sommer, R. J. (2020). Phenotypic plasticity: From theory and genetics to current and future challenges. Genetics, 215(1):1–13.

Svardal, H., Rueffler, C., and Hermisson, J. (2011). Comparing environmental and genetic variance as adaptive response to fluctuating selection. Evolution, 65(9):2492–2513.

Svardal, H., Rueffler, C., and Hermisson, J. (2015). A general condition for adaptive genetic polymor-phism in temporally and spatially heterogeneous environments. Theoretical Population Biology, 99:76–97.

Sæther, B.-E. and Engen, S. (2015). The concept of fitness in fluctuating environments. Trends in Ecology and Evolution, 30(5):273–281.

Tielbörger, K., Bilton, M. C., Metz, J., Kigel, J., Holzapfel, C., Lebrija-Trejos, E., Konsens, I., Parag, H. A., and Sternberg, M. (2014). Middle-Eastern plant communities tolerate 9 years of drought in a multi-site climate manipulation experiment. Nature Communications, 5(1).

Tielbörger, K. and Petrů, M. (2010). An experimental test for effects of the maternal environment on delayed germination. Journal of Ecology, 98(5):1216–1223.

Tielbörger, K. and Prasse, R. (2009). Do seeds sense each other? Testing for density-dependent germination in desert perennial plants. Oikos, 118(5):792–800.

Tielbörger, K. and Valleriani, A. (2005). Can seeds predict their future? Germination strategies of density-regulated desert annuals. Oikos, 111(2):235–244.

Tufto, J. (2000). The evolution of plasticity and nonplastic spatial and temporal adaptations in the presence of imperfect environmental cues. The American Naturalist, 156(2):121.

Turcotte, M. M. and Levine, J. M. (2016). Phenotypic plasticity and species coexistence. Trends in Ecology & Evolution, 31(10):803–813.

Venable, D. L. and Lawlor, L. (1980). Delayed germination and dispersal in desert annuals: Escape in space and time. Oecologia, 46(2):272–282.

Venable, D. L., Pake, C. E., and Caprio, A. C. (1993). Diversity and coexistence of Sonoran desert winter annuals. Plant Species Biology, 8(2-3):207–216.

Via, S., Gomulkiewicz, R., De Jong, G., Scheiner, S. M., Schlichting, C. D., and Van Tienderen, P. H. (1995). Adaptive phenotypic plasticity: Consensus and controversy. Trends in Ecology and Evolution, 10(5):212–217.

Via, S. and Lande, R. (1985). Genotype-environment interaction and the evolution of phenotypic plasticity. Evolution, 39(3):505–522.

Visser, M. E. and Gienapp, P. (2019). Evolutionary and demographic consequences of phenological mismatches. Nature Ecology & Evolution, 3:879–885.

West-Eberhard, M. J. (1989). Phenotypic plasticity and the origins of diversity. Annual Review of Ecology and Systematics, 20(1):249–278.

Wisnoski, N. I. and Shoemaker, L. G. (2022). Seed banks alter metacommunity diversity: The interactive effects of competition, dispersal and dormancy. Ecology Letters, 25(4):740–753.

Woltereck, R. (1909). Weitere experimentelle Untersuchungen über Artveränderung, speziell über das Wesen quantitativer Artunterschiede bei Daphnien. Verhandlungen der deutschen zoologischen Gesellschaft, 19:110–173.

Yamamichi, M. and Hoso, M. (2017). Roles of maternal effects in maintaining genetic variation: Maternal storage effect. Evolution, 71(2):449–457.

Yamamichi, M., Jr, N. G. H., Rees, M., and Ellner, S. P. (2019). Rapid evolution with generation overlap : The double-edged effect of dormancy. Theoretical Ecology, 12(2):179–195.

Yamamichi, M., Letten, A. D., and Schreiber, S. J. (2023). Eco-evolutionary maintenance of diversity in fluctuating environments. Ecology Letters, 26(S1):S152–S167.

